# A PTBP1–CDC42 splicing axis regulates leukemia growth and venetoclax sensitivity in acute myeloid leukemia

**DOI:** 10.64898/2026.08.25.745954

**Authors:** Margaux Oberling, Mathieu Landry, Yann Aubert, Manon Faivre, Alexandre Gay, Alexandre Boudet, Anaïs Granjon, Ambrine Sahal, Sarah Bertoli, François Vergez, Véronique Mansat-De Mas, Christian Récher, Clément Larrue, Laura Poillet, Jean-Emmanuel Sarry, Carine Joffre, Manuel D. Diaz-Munoz, Vera Pancaldi, Margherita Ghisi

## Abstract

Acute myeloid leukemia (AML) is an aggressive blood cancer characterized by high rates of relapse and poor outcomes, especially in elderly or unfit patients, who cannot tolerate intensive chemotherapy. While the BCL2 inhibitor venetoclax has improved initial responses in this high-risk population, relapses remain nearly universal, highlighting the need for novel therapeutic strategies. Here, we identify the RNA-binding protein PTBP1 as a critical dependency in AML. PTBP1 depletion impairs leukemic growth *in vitro* and *in vivo*, and is associated with widespread splicing alterations and global disruption of protein synthesis. Integrative transcriptomic and iCLIP analyses reveal that PTBP1 orchestrates a splicing program centered on Rho GTPase signaling, with CDC42 as a key downstream effector. Mechanistically, PTBP1 loss triggers a splicing switch from CDC42-v1 to CDC42-v2, leading to reduced GTPase activity and impaired protein synthesis. Pharmacological inhibition of CDC42 selectively induces cytotoxicity in AML cells, while sparing healthy hematopoietic cells. Importantly, CDC42 inhibition markedly enhances venetoclax anti-leukemic efficacy. These findings establish PTBP1 as a critical regulator of AML cell fitness and identify a clinically actionable therapeutic combination that exploits AML dependency on PTBP1-CDC42 signaling to enhance the efficacy of venetoclax-based regimens.

## INTRODUCTION

Acute myeloid leukemia (AML) is the most common and lethal form of leukemia in adults, with an average 5-year overall survival rate of less than 30%, dropping to 10% in elderly patients. This poor prognosis is mainly due to high rates of relapse after chemotherapy, driven by persistent drug-tolerant leukemic subpopulations(1–3). Outcomes are particularly poor for elderly and medically unfit patients who are ineligible for intensive chemotherapy and have limited therapeutic options. The recent introduction of combination regimens based on the BCL2 inhibitor venetoclax as standard of care for these patients has significantly improved initial response rates; however, responses are typically transient and almost invariably followed by relapse(4). This clinical context highlights the urgent need to deepen our understanding of AML biology and to identify novel, actionable molecular vulnerabilities and therapeutic combinations to improve patient survival.

Various mechanisms have been proposed to contribute to venetoclax resistance, depicting a highly complex scenario involving both mutation-dependent and mutation-independent mechanisms. These include mutations in the BCL-2 binding groove that reduce drug affinity, compensatory upregulation of anti-apoptotic proteins such as MCL-1 or BCL-XL, alterations in energy metabolism or specific signaling pathways, all of which have been associated with poor responses to venetoclax regimens(5). Additionally, emerging evidence suggests that targeting RNA splicing machinery(6,7) or protein synthesis(8,9) may represent promising strategies to overcome venetoclax resistance.

RNA splicing is an essential step in mRNA maturation and gene expression regulation, enabling the generation of multiple transcript isoforms with distinct functions and coding potential from a single gene. Aberrant RNA splicing has emerged as a key contributor to cancer pathogenesis(10). In AML, interest in the role of splicing defects in leukemia initiation and progression has intensified following the identification of recurrent mutations in spliceosome components(11–13). However, a series of studies in patients has shown that splicing alterations are widespread in AML even in the absence of mutations affecting the splicing machinery, with up to one-third of expressed genes being differentially spliced in leukemic cells compared with healthy CD34⁺ progenitors(14). Notably, this extensive splicing rewiring affects leukemia-associated genes(15), and has been shown to drive AML leukemogenesis(16,17), survival(18) and therapy evasion(6,19). Collectively, these findings indicate that leukemic cells recurrently rely on altered splicing programs, creating a potential non-oncogene addiction to specific splicing regulators that may be therapeutically exploited.

Polypyrimidine tract binding protein 1 (PTBP1) is a multifunctional RNA-binding protein that regulates mRNA splicing, translation, stability, and subcellular localization(20). While PTBP1 is best characterized for its role in neuronal lineage commitment(21), its dysregulation has also been implicated in tumorigenesis and associated with poor prognosis across multiple cancers(20,22,23). Emerging evidence now suggests a critical but incompletely understood role for PTBP1 in normal and neoplastic hematopoietic cells. In murine models, PTBP1 was shown to regulate key aspects of normal hematopoiesis, including hematopoietic stem cell maintenance, erythropoiesis, and B-cell selection(24,25). In addition, PTBP1 has been implicated in the regulation of autophagy, metabolism, and cell survival in specific AML subtypes(26–29).

In this work, we demonstrate that PTBP1 is essential for AML growth and survival, both *in vitro* and *in vivo*. Our results suggest that this dependency is independent of basal PTBP1 expression levels or mutational background, and is associated with widespread alteration of RNA splicing and global impairment of protein synthesis upon PTBP1 depletion. Analysis across multiple AML models identifies a conserved PTBP1-dependent splicing program and implicates Rho GTPase signaling as a key, actionable pathway downstream of PTBP1. Consistent with this model, pharmacological inhibition of the Rho GTPase CDC42 induces protein synthesis inhibition and markedly enhances the efficacy of venetoclax in AML cells.

## MATERIAL AND METHODS

### AML Cell lines and primary samples

The human AML cell lines MV4-11 and OCI-AML3 were obtained from DSMZ (Leibniz Institute, Germany), MOLM-14 from M. Carroll (University of Pennsylvania, Philadelphia, USA), and U937 from ATCC (American Type Culture Collection, Manassa, VA, USA). Mutational features of these AML cell lines are detailed in Table S1.

Primary AML cells were collected from peripheral blood samples during routine diagnostic procedures at Toulouse University Hospital (TUH) after obtaining informed consent from patients. The clinical and biological annotations of the samples were declared to the CNIL (Comité National Informatique et Libertés; ‘Data processing and Liberties National Committee’). The cytogenetic and genomic characteristics of the leukemic samples (TUH) used in this study are described in Table S2.

Human peripheral blood mononuclear cells (PBMCs) were isolated from freshly-drawn blood from healthy donors. All human samples were collected after informed consent and under an approved protocol.

Purification details for primary AML cells and PMBCs, as well as cell culture conditions are provided in the Supplemental Information.

### Lentiviral-mediated shRNA knockdown

For lentiviral production, HEK293T/17 cells were co-transfected with the lentiviral packaging constructs p8.91 (4 μg; Addgene plasmid #187441), pVSV-G (2 μg; Addgene plasmid #138479), and 6 μg of the shPTBP1 or shCTL (shRNA against Renilla Luciferase) plasmid to generate lentivirus. SGEN miR-E-based RNAi plasmids (Addgene plasmid #111171) were used for constitutive shRNA expression, and LT3GEPIR Tet-ON miR-E-based RNAi plasmids (Addgene plasmid #111177) were used for doxycycline-inducible shRNA expression(30). In both vectors the expression of the shRNA is linked to the expression of the reporter GFP. P8.91 and pVSV-G plasmids were a gift from Simon Davis and Akitsu Hotta, respectively. The SGEN and LT3GEPIR vectors (a gift from Johannes Zuber, IMP, Vienna) were modified to insert shRNA sequences targeting PTBP1. ShRNA sequences are provided in the Supplemental Information. Cloning procedure and transduction protocol for leukemia cells are available on request.

### AML mouse xenograft model

All animal procedures were approved by the Institutional Animal Care and Use Committee of Region Midi-Pyrenees and the Ministry of Higer Education and Research (France, Apafis N° 32669-2021080215068127 v4, 52276-2024112714182243 v2). NSG (NOD/LtSz-SCID/IL-2Rα-/-) mice were produced at the Genotoul Anexplo platform in Toulouse (France) using breeders obtained from Charles River Laboratories. Mice were housed in sterile conditions using HEPA-filtered microisolators and fed with irradiated food and sterile water. For CLDX generation, NSG mice (6–10 weeks old, male/female) were sub-lethally conditioned with busulfan (20 mg/kg, i.p.), and, after 48 h, injected *via* the tail vein with leukemic cells (0.2-2×10^6^ cells/100µL in HBSS). Mice were randomly assigned to experimental groups based on sex and body weight. The animals were daily monitored for disease symptoms, and humanly sacrificed upon the appearance of signs of distress, in accordance with ethical guidelines. To assess leukemic engraftment, the mice were humanely euthanized in accordance with European ethics protocols at day 18 post-injection. Bone marrow (from tibias, femurs, and hips) and spleen were harvested, mechanically dissociated in HBSS with 2%FBS, and processed into single-cell suspensions for flow cytometric analysis and/or FACS sorting of leukemic cells. Following RBC lysis, human leukemic cells were labeled with fluorochrome-conjugated hCD45 antibodies and AnnexinV to determine the fraction of viable transduced human blasts (AnxV-/hCD45+/GFP+).

### Flow cytometry-based assays

Viability was assessed using flow cytometry with AnnexinV (BD horizon) and Propidium Iodide (PI, Sigma Aldrich) staining.

Global protein synthesis was measured by cytofluorimetric evaluation of puromycin incorporation. AML cells were incubated with 10 µg/mL puromycin (InvivoGen) for 9 minutes, washed with ice-cold PBS, and fixed/permeabilized with FoxP3 Fix/Perm Buffer (20 minutes incubation at room temperature; eBioscience, Thermo Fisher Scientific). Cells were then incubated overnight at 4°C with an anti-puromycin antibody (BioLegend), and puromycin labeling mean fluorescence intensity (MFI) was quantified the following day.

All flow cytometry data were acquired using a CytoFLEX flow cytometer (Beckman Coulter), and analyzed in FlowJo (Becton Dickinson).

### Cell sorting

Leukemic cells were stained with fluorochrome-conjugated antibodies targeting hCD45, hCD33 or hCD44, along with a viability stain. Sorting was performed on a BD FACS Aria Fusion and FACS Melody Sorters. Details on antibodies and reagents are provided in the Supplemental Information.

### Colony Forming Assay

For colony forming assays, 1 000 FACS-sorted transduced AML cells (hCD45+/GFP+/AnxV-) were plated in duplicate in 1 mL of methylcellulose medium (H4230, Stemcell Technologies), and colonies were counted 7-10 days later by light microscopy upon staining with MTT (Thiazolyl blue tetrazolium bromide).

### Global and targeted analysis of small GTPase signaling and pan-kinase activity

Global Serine/Threonine (STK) and Tyrosine kinase (PTK) activity, was measured using a PamGene Kinase Activity Assay with a PamChip® peptide microarray and a PamStation12 workstation (Pamgene). This assay was performed on MOLM-14, MV4-11, and U937 cells constitutively expressing shPTBP1 or shCTL, 3 days after transduction with shRNA vectors. The Median Kinase Statistic represents the median fold-change (Log_10_) in kinase activity relative to the control, indicating significant changes in activity between shPTBP1 and shCTL conditions across all three cell lines.

Total GTPase activity and CDC42 activity were quantified using the GTPase-Glo^TM^ Assay (Promega, Cat#V7681) and the Cdc42 G-LISA® GTPase Activation Assay (Cytoskeleton Inc., Cat#BK127), respectively, following the manufacturer’s instructions.

### RNA-seq

Viable transduced (AnxV-/GFP+) AML cells (MOLM-14, MV4-11, OCI-AML3, U937) were FACS-sorted two days after transduction with SGEN shPTBP1 (#1 and #2) or shCTL lentiviral vectors. Three days after transduction, total RNA was isolated using the RNeasy Plus Micro kit with gDNA removal (Qiagen), according to the manufacturer’s instructions. RNA quantification, purity and integrity was monitored using NanoDrop/Qubit quantification/integrity measurement. Only RNA with no sign of contamination or marked degradation (RQN>9) were used for further analysis. The generation and sequencing of cDNA libraries was performed by the LuxGen genomic platform (Luxembourg Institute of Health, Luxembourg). PolyA-selected libraries were prepared using the TruSeq® Stranded mRNA Library Prep Kit (Illumina) and subjected to paired-end (150 bp) sequencing on the NextSeq 550 System (Illumina) with a depth of 100M reads/sample. Two biological replicates per condition per cell line were made.

### Individual Cross-Linking Immunoprecipitation (iCLIP)

PTBP1:RNA interactome in MOLM-14 cells was annotated using individual cross-linking immunoprecipitation (iCLIP), as previously described(31,32). The detailed protocol is available in the Supplemental Information.

### Bioinformatics analysis

Differential gene expression analysis was conducted using DESeq2 (v1.44.0), using default parameters, with CombatSeq used to adjust for batch effects. Significant changes in gene expression were defined as those with a Benjamini-Hochberg adjusted p-value (padj) ≤ 0.05. Alternative splicing analysis was performed with rMATS turbo (v4.1.2), with Maser filtering (F.T. Veiga D (2025). *Maser: Mapping Alternative Splicing Events to pRoteins*. R package version 1.29.0, https://bioconductor.org/packages/maser). Differential Splicing events were filtered for those with an inclusion level difference (deltaPSI) > 10%, an FDR < 0.05, and average reads >10. The functional consequences of PTBP1 depletion on isoform switching were analyzed using IsoformSwitchAnalyzeR (v1.18.0).

Gene list functional enrichment analysis was performed using EnrichR or the gProfiler2 R package, filtering out enrichment terms defined by more than 1500 genes and by less than 15 genes, as indicated.

To define PTBP1:RNA interactome, iCLIP data was mapped to the human genome GRCh38/hg38 and analyzed using the iMaps pipeline(33,34) in Flow (https://app.flow.bio/). PTBP1 crosslink sites were filtered for those with an FDR ≤ 0.05 and an extended_score ≥ 3. Genomic distribution of PTBP1 crosslink sites was annotated using ChIPSeeker with TxDb.Hsapiens.UCSC.hg38.known Gene annotation. Direct PTBP1 targets were defined by integrating iCLIP and RNA-seq analysis on shPTBP1 vs. shCTL AML cells, as detailed in the Supplemental Information.

PTBP1 target protein-protein interaction network was generated using String v.12.0 (https://string-db.org/).

### Statistical Analysis

All statistical analyses were conducted using GraphPad Prism software v9.2.0 (RRID: SCR_002798). Statistical significance was assessed using two-tailed unpaired or paired Student’s t test, with Welch correction applied when variances were unequal. Kaplan-Meier analysis was used for survival data. Sample sizes (N) are specified in the figure legends. Data are presented as mean ± standard deviation. *P* values ≤ 0.05 were considered to be significant (*, P < 0.05; **, P < 0.01; ***, P < 0.001; ****, P < 0.0001).

## RESULTS

### AML cells depend on the splicing factor PTBP1 for their growth and survival

Analysis of publicly available transcriptomic datasets showed that *PTBP1* transcript levels vary widely across AML patients and did not correlate with overall survival (Fig. S1A-B). Similarly, analysis of PTBP1 protein abundance in a limited panel of primary AML samples and established cell lines confirmed substantial heterogeneity, with no clear association with recurrent genetic lesions (Fig. S1C-D; Table S1, 2).

To investigate the functional role of PTBP1 in AML, a panel of genetically diverse human AML cell lines (MOLM-14, MV4-11, OCI-AML3 and U937) were transduced with lentiviral vectors expressing PTBP1-targeting shRNAs (shPTBP1) or a non-targeting control (shCTL), coupled to GFP expression (Fig. 1A-B; Fig.S1E). PTBP1 knock-down (KD) resulted in a marked negative selection of PTBP1-depleted cells in competitive growth assays (Fig. 1C), irrespective of their mutational background or basal PTBP1 expression levels (Fig. S1C; Table S1). This phenotype was confirmed using a second independent shRNA, ruling out off-target effects (Fig. S1F-G). Further investigations demonstrated that PTBP1 KD impaired leukemic growth in liquid culture (Fig. 1D; Fig. S1H) and reduced clonogenic capacity in methylcellulose assays (Figure 1E-F; Fig. S1I). This growth defect was associated with a modest and delayed induction of apoptosis, detectable from day 6 post-transduction (Fig. 1G).

**Figure 1.**
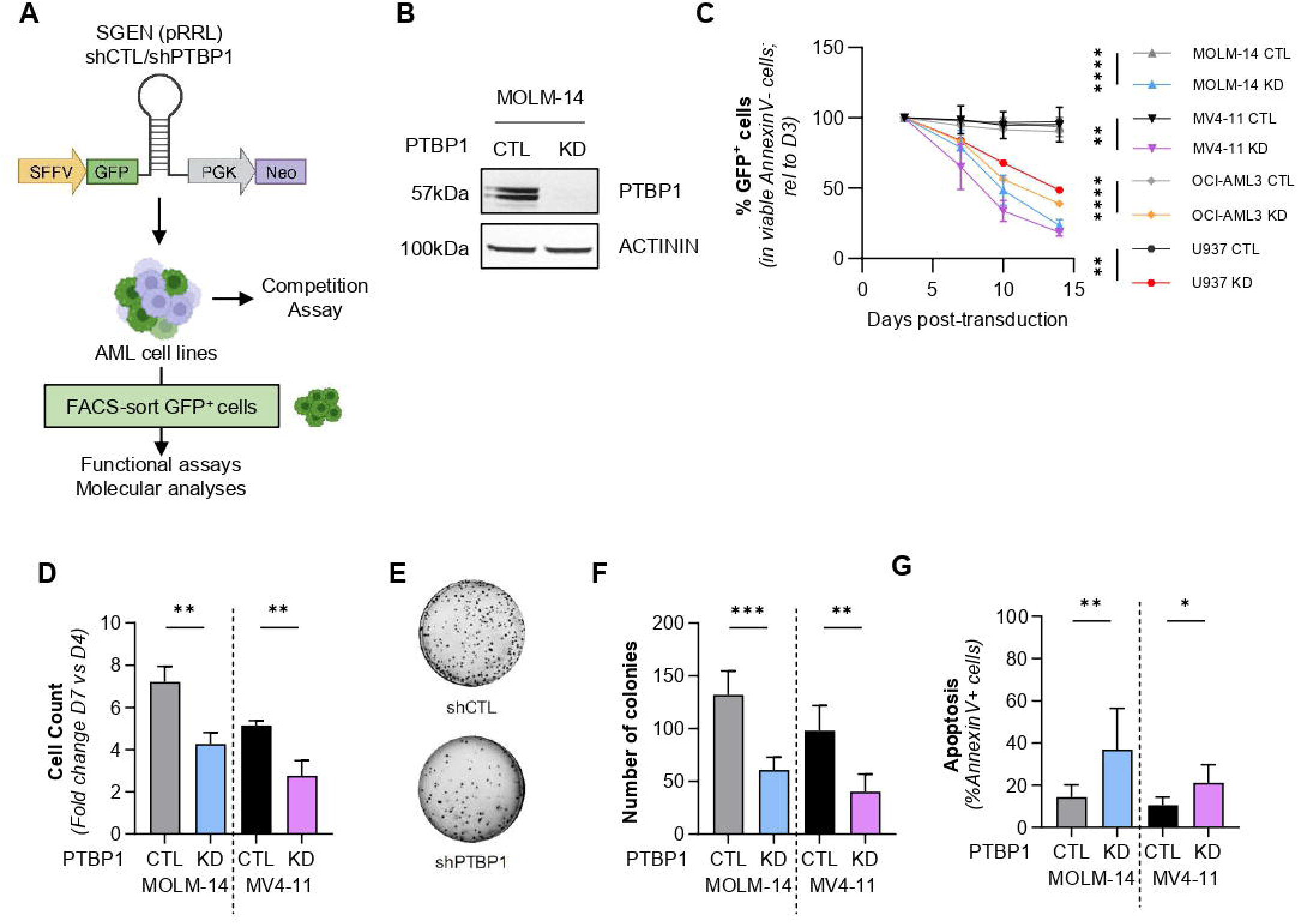
PTBP1 depletion impairs leukemic growth of AML cells *in vitro*. **(A)** Schematic overview of PTBP1 knock-down (KD) experimental strategy and MiRE-based shRNA-expressing lentiviral vector (SGEN-shRNA). PTBP1 silencing was achieved by transduction of AML cell lines (MOLM-14, MV4-11, OCI-AML3 and U937) with a MiRE-based lentiviral vector (SGEN backbone;(30)) constitutively expressing an shRNA against PTBP1 (shPTBP1) linked to the fluorescent reporter GFP. An shRNA targeting the *Renilla luciferase* gene was used as negative control (shCTL). **(B)** Western Blot analysis of PTBP1 expression in GFP-positive FACS-sorted AML cells 6 days after transduction with SGEN-shPTBP1(PTBP1 KD) or -shCTL (CTL) lentiviral vectors. ACTININ was used as a loading control. **(C)** Competitive proliferation assay on mixed AML population including GFP-positive (shRNA-expressing) and GFP-negative (untransduced) cells. The percentage of viable GFP-positive cells was monitored over time by flow cytometry and normalized to day 3. Data are presented as mean values ± SD. N=3. **(D)** Trypan blue staining-based cell counting of PTBP1 KD *vs.* CTL AML cells. Viable shRNA-expressing (PI^-^/GFP^+^) AML cells were FACS sorted 3 days after transduction and plated at the same concentration. Viable cell counts at day 7 were normalized by input counts at day 4. N=3 **(E-F)** Methylcellulose colony forming assay on GFP-positive, shRNA-expressing AML cells. FACS-sorted GFP-positive cells were plated in semi-solid medium 3 days after transduction. The number of colonies was assessed by MTT staining 10 days after plating. A representative image of MOLM-14 cells **(E)** and cumulative data for MOLM-14 and MV4-11 cell lines **(F)** and are shown. N=5 for MOLM-14, N=4 for MV4-11. **(G)** Flow cytometric analysis of apoptosis by Annexin V staining. The percentage of apoptotic (Annexin V-positive) cells was measured in GFP-positive (shRNA-expressing) cells 6 days after transduction. N=9 for MOLM-14, N=5 for MV4-11.

We next examined the impact of PTBP1 depletion on leukemia growth and maintenance *in vivo*. GFP-positive MOLM-14 and MV4-11 human AML cell lines transduced to express shPTBP1 or shCTL were FACS-purified and injected intravenously into immunodeficient NSG (NOD-scid IL2Rgamma ^null^) mice (Fig. 2A; Fig. S2A). PTBP1 KD significantly prolonged survival of xenografted mice compared to controls (Fig. 2B). Flow cytometric analysis of bone marrow performed 18 days post-engraftment revealed a severe reduction in leukemic burden in mice injected with PTBP1-depleted AML cells, both in terms of blast frequency and absolute cell number (Fig. 2C-D). Of note, despite injection of leukemic cells purified to >95% GFP⁺, we observed a strong negative selection against shPTBP1-expressing blasts *in vivo* (Fig. 2E), accompanied by increased apoptosis in the residual PTBP1-depleted leukemic population (Fig. 2F).

**Figure 2.**
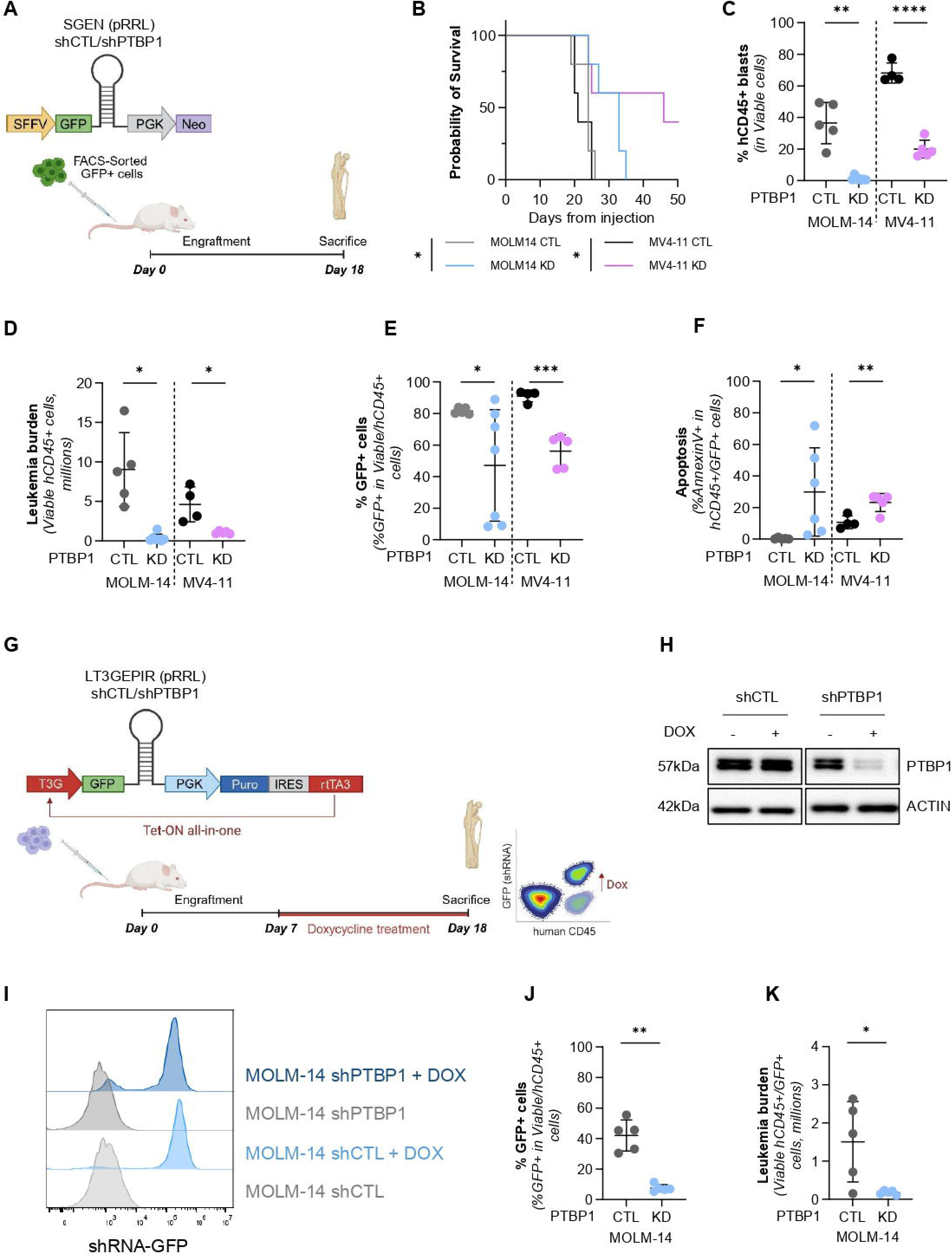
PTBP1 depletion impairs leukemia engraftment and growth in preclinical models of AML. **(A)** Schematic overview of the PTBP1 knock-down (KD) experimental strategy *in vivo.* MOLM-14 cells and MV4-11 AML cells were transduced with SGEN-shRNA lentiviral vectors (30) constitutively expressing an shRNA against PTBP1 (shPTBP1) or a negative control targeting *Renilla luciferase* (shCTL), both linked to the fluorescent reporter GFP. Viable (AnxV-) GFP-positive, shRNA-expressing cells were FACS-sorted 3 days after transduction and engrafted via intravenous injection into immunodeficient NSG mice. Mice were sacrificed at day 18 post-engraftment to evaluate leukemia burden in the bone marrow. **(B)** Kaplan-Meier survival curve of mice injected with PTBP1 KD or CTL MOLM-14 and MV4-11 AML cells. N=5 mice per group. **(C-F)** Cytofluorimetric analysis of AML cells harvested from the bone marrow of mice 18 days after leukemia injection. **(C-D)** Percentage and absolute number of viable CD45-positive human AML blasts. **(E)** Percentage of GFP-positive (shRNA-expressing) cells within the viable human AML population. **(F)** Apoptosis quantification in CD45-positive/ GFP-positive human AML cells assessed by cytofluorimetric analysis upon AnnexinV staining. Data are presented as mean values ± SD of biological replicates. **(G)** Schematic overview of the doxycycline inducible PTBP1 knockdown (KD) experimental strategy *in vivo*. MOLM-14 cells were transduced with LT3GEPIR lentiviral vectors (30) expressing an shRNA against PTBP1 (shPTBP1) or a negative control targeting *Renilla luciferase* (shCTL), both linked to the fluorescent reporter GFP under the T3G Tet-responsive promoter. The vector also included a puromycin-resistance cassette and the Tet-On transcriptional activator (rtTA) under a constitutive promoter, enabling doxycycline-inducible shRNA and GFP expression *in vitro* and *in vivo*. After puromycin selection, transduced MOLM-14 cells were injected intravenously into NSG mice. One week after injection, doxycycline was administered *via* drinking water to induce shRNA expression. Mice were sacrificed at day 18 post-engraftment to evaluate leukemia burden in the bone marrow. **(H)** Western blot analysis of PTBP1 expression in MOLM-14 cells transduced with LT3GEPIR lentiviral vectors expressing shRNAs targeting PTBP1 (shPTBP1#1) or a control shRNA (shCTL) after 3 days of doxycycline treatment (+/-) *in vitro*. β-ACTIN was used as a loading control. **(I)** Representative flow-cytometry profiles showing GFP expression switch in MOLM-14 cells transduced with LT3GEPIR-shPTBP1 lentiviral vector following doxycycline treatment (+/-). **(J-K)** Cytofluorimetric analysis of AML cells harvested from the bone marrow of mice 18 days after injection of LT3GEPIR-transduced leukemia. Percentage **(J)** and absolute number **(K)** of viable GFP-positive (shRNA-expressing) CD45-positive human AML blasts in the bone marrow of recipient mice.

To assess whether PTBP1 is also required for leukemia maintenance, we established MOLM-14 xenografts expressing doxycycline-inducible shRNAs targeting PTBP1 or a negative control, linked to a GFP reporter (Fig. 2G-I). Following leukemia establishment, PTBP1 depletion was induced by administering doxycycline in the drinking water of the animals. Strikingly, we observed a marked decrease in both the percentage and absolute number of shPTBP1-expressing leukemic cells (GFP-positive) in the bone marrow (Fig. 2J-K), indicating that PTBP1 is not only required for leukemia engraftment but also for its ongoing maintenance *in vivo*.

Collectively, our findings establish PTBP1 as a critical dependency in human AML and underscore the need to explore the downstream splicing programs and molecular pathways that mediate its critical role in leukemia.

### PTBP1 depletion drives widespread splicing alterations in AML cells

It has been proposed that PTBP1 might modulate glucose metabolism in both physiological and cancer contexts by controlling alternative splicing of the glycolytic enzyme pyruvate kinase M (PKM). By repressing the PKM1 isoform, PTBP1 can lower the PKM1/PKM2 ratio, which has been shown to promote a metabolic shift from mitochondrial oxidative phosphorylation to anaerobic glycolysis, a hallmark of the Warburg Effect in highly anabolic and rapidly proliferating cells, such as cancer cells(35,36). In other cancer models, PTBP1’s oncogenic function has been linked to its role in the control of mitochondrial metabolism. We therefore investigated whether the growth defect induced by PTBP1 depletion was associated with its metabolic function. Although PTBP1 knockdown in AML cells increased PKM1 expression and PKM1/PKM2 ratio (Fig. S3A), its effects on glycolytic and oxidative metabolism - assessed by oxygen consumption rate (OCR), extracellular acidification rate (ECAR), and mitochondrial ATP production quantification - were highly heterogeneous across AML models (Fig. S3B-G). These results strongly suggest that the growth inhibitory effects of PTBP1 depletion in human AML cells are independent of alterations in glycolytic or oxidative metabolic states.

To investigate how PTBP1 sustains AML cell growth at the molecular level, we performed RNA sequencing (RNA-seq) on four human AML cell lines (MOLM-14, MV4-11, OCI-AML3 and U937) following PTBP1 KD. This analysis revealed relatively modest transcriptional alterations with only 289, 137, 118 and 150 genes significantly differentially expressed (FDR ≤ 0.05, |log₂FC| ≥ 1) upon PTBP1 KD in MOLM-14, MV4-11, OCI-AML3 and U937 cells, respectively (Fig. 3A-B). Cross-comparison of these gene lists identified only 24 genes (of which only 16 were annotated) commonly deregulated across all AML models, including *PTBP1* and its paralog *PTBP2*, underscoring the largely cell line–specific effects of PTBP1 depletion on mRNA abundance (Fig. 3C-D). To assess whether these transcriptional changes converged on shared biological processes, we performed comparative over-representation analysis (ORA) of the differentially expressed genes in PTBP1 KD *vs.* CTL condition in each cell line. This analysis confirmed substantial heterogeneity in PTBP1-regulated pathways, with a modest enrichment for genes involved in lipid and carbohydrate derivative metabolism (Fig. S4A).

**Figure 3.**
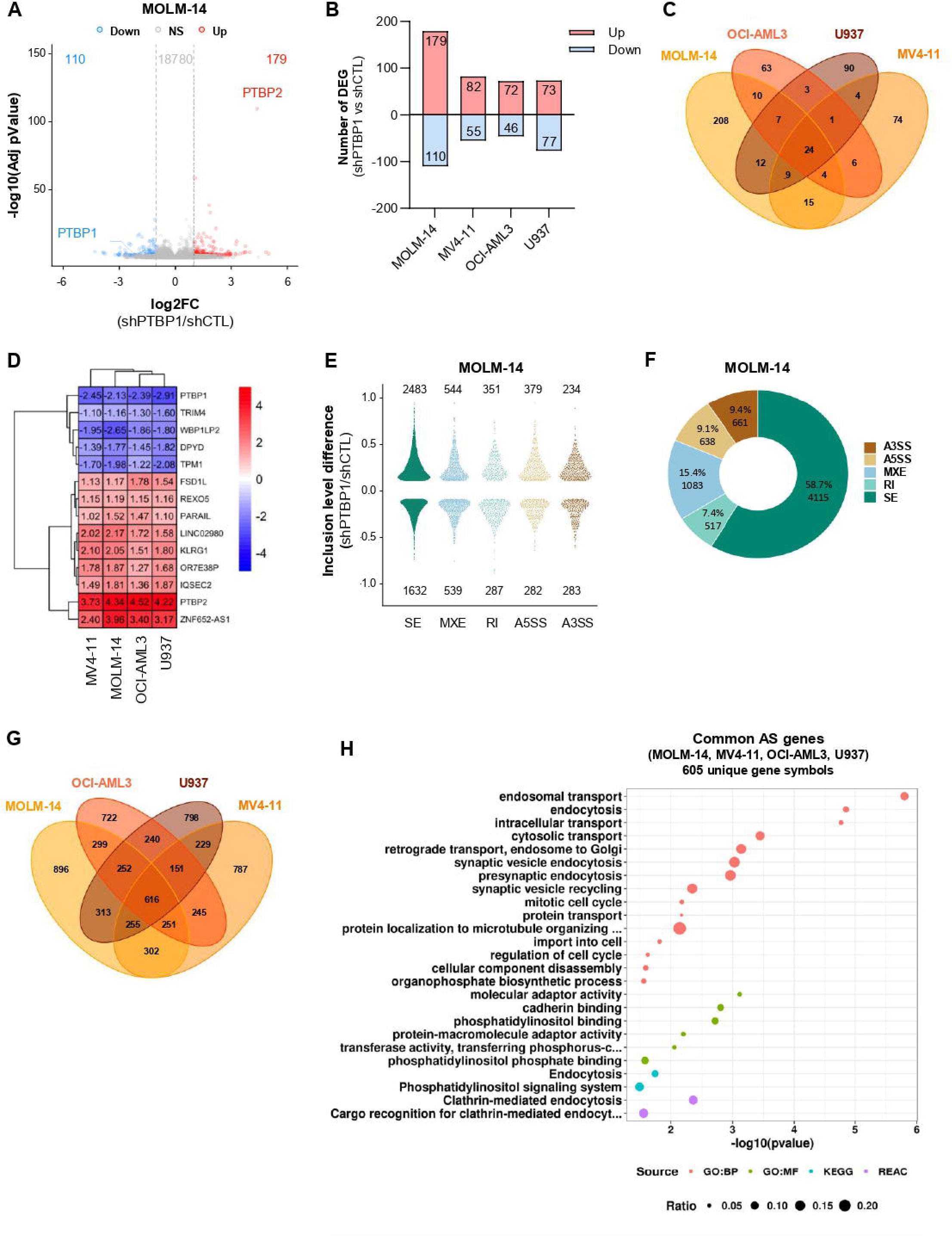
PTBP1-mediated regulation of transcript abundance and alternative splicing in human AML cells. RNA-seq analyses were performed on GFP-positive AML cells FACS-sorted 3 days after transduction with lentiviral vectors expressing PTBP1-targeting shRNAs (shPTBP1) or a negative control shRNA (shCTL). Alternative splicing (AS) and differential gene expression analyses were then performed as described in the Methods. **(A)** Volcano plot representing differential gene expression in shPTBP1 compared to shCTL MOLM-14 AML cells. Colored dots indicate genes with absolute log2 fold change (FC) > 1 (red) and < 1 (blue) and adjusted p-value ≤ 0.05. **(B)** Barplot showing the number of differentially expressed genes (DEG) (absolute log2 FC ≥ 1, FDR ≤ 0.05) in the shPTBP1 *vs.* shCTL condition in the indicated AML cell lines. **(C)** Venn diagram illustrating the overlap of gene sets differentially expressed in the shPTBP1 *vs.* shCTL condition across different AML cell lines. **(D)** Heatmap illustrating DEGs commonly regulated upon PTBP1 KD across the 4 AML cell lines tested. Of the 24 genes identified, only 16 are annotated genes and are shown. Blue indicates negative FC (log2 FC), corresponding to lower expression in PTBP1 KD cells *vs.* CTL; red indicates positive FC (log2 FC), corresponding to higher expression in PTBP1 KD cells *vs.* CTL. **(E)** Beanplot representing the distribution of alternative splicing (AS) events inclusion level difference in PTBP1 KD *vs.* CTL MOLM-14, binned by AS event type (SE: Skipped Exon, MXE: Mutually Exclusive Exon, RI: Retained Intron, A5SS and A3SS: Alternative 5’ or 3’ Splice Site). AS events with Incl. Level. Diff ≥ 0.1, FDR ≤ 0.05, avg. read counts>10 are shown and numbers are indicated on the plot. Differential AS events were identified by rMATS and filtered using maser R package. **(F)** Donut chart illustrating the distribution of differential AS events across specific AS event types in PTBP1 KD *vs.* CTL MOLM-14 cells. Both the absolute number and relative proportion of each AS event type are shown. **(G)** Venn diagram representing the overlap between differentially AS genes identified in the PTBP1 KD *vs.* CTL condition across the 4 AML cell lines tested. **(H)** Pathway enrichment analysis of the differentially AS genes shared across all 4 AML cell lines, identified through comparison of PTBP1 KD and CTL AS gene sets. Of the 616 genes identified, only 605 corresponding to unique gene symbols were considered for the analysis. GProfiler and the indicated source databases (GO:BP=Gene Ontology:Biological Processes; GO:MF=Gene Ontology:Molecular Functions; KEGG; REAC=Reactome) were used for the analysis.

In stark contrast, alternative splicing (AS) analysis using rMATS (FDR ≤ 0.05, |Incl. Level Diff.| > 10%, average read count > 10) revealed widespread splicing deregulation following PTBP1 KD, affecting over 2 500 genes per cell line (MOLM-14: 3 075; MV4-11: 2 706; OCI-AML3: 2 674; U937: 2 752) (Fig. 2E; Fig. S4B).

Consistent with its established role as a context-dependent splicing regulator controlling alternative exon inclusion/exclusion, exon skipping (SE) represented the predominant AS event affected by PTBP1 perturbation (>50% of differentially spliced events), followed by mutually exclusive exon inclusion (MXE; ∼15%) (Fig. 2F; Fig. S4C). PTBP1 KD generally resulted in increased exon inclusion, indicating that PTBP1 primarily acts as a splicing repressor in AML cells. Notably, only a small fraction of differentially spliced genes showed concomitant changes in mRNA abundance, suggesting largely independent regulation of splicing and transcript abundance (Fig. S4D).

Across all four AML cell lines, we identified 616 genes whose splicing was consistently altered upon PTBP1 depletion (Fig. 3G). Pathway enrichment analysis of this shared splicing signature revealed a significant over-representation of genes involved in endocytosis, intracellular vesicle transport, and cell cycle regulation (Fig. 3H). Beyond this common signature, comparative over-representation analysis of the top 500 alternatively spliced genes in each cell line confirmed a shared and highly conserved enrichment of genes governing these processes (Fig. S4E).

These findings highlight PTBP1 as a dominant and conserved regulator of alternative splicing in AML cells, with his loss leading to profound and widespread splicing alterations that converge on key cellular processes.

### PTBP1 regulates a Rho GTPase–centered splicing program that sustains AML protein synthesis and cell proliferation

To identify direct PTBP1 splicing targets and gain mechanistic insight into its mode of action, we performed iCLIP (individual nucleotide-resolution UV Cross-Linking and Immuno-Precipitation) in MOLM-14 cells (Fig. S5A–C). This analysis revealed that over 90% of PTBP1 binding sites were localized within intronic regions (Fig. 4A), consistent with a primary role in splicing regulation rather than in mRNA stability or translation in AML cells. Integration of iCLIP and RNA-seq data identified alternative splicing events directly regulated by PTBP1, defined by PTBP1 binding proximal to regions that were differentially spliced upon PTBP1 KD (Fig. S5D). The proportion of AS events bound by PTBP1 varied by splicing event category, with 40–50% of SE and MXE events showing proximal PTBP1 occupancy, compared with markedly lower proportions for alternative 3′ and 5’ splice site (A3SS: 20%, A5SS: 3%), and intron retention (RI: 21%) events (Fig. 4B).

**Figure 4.**
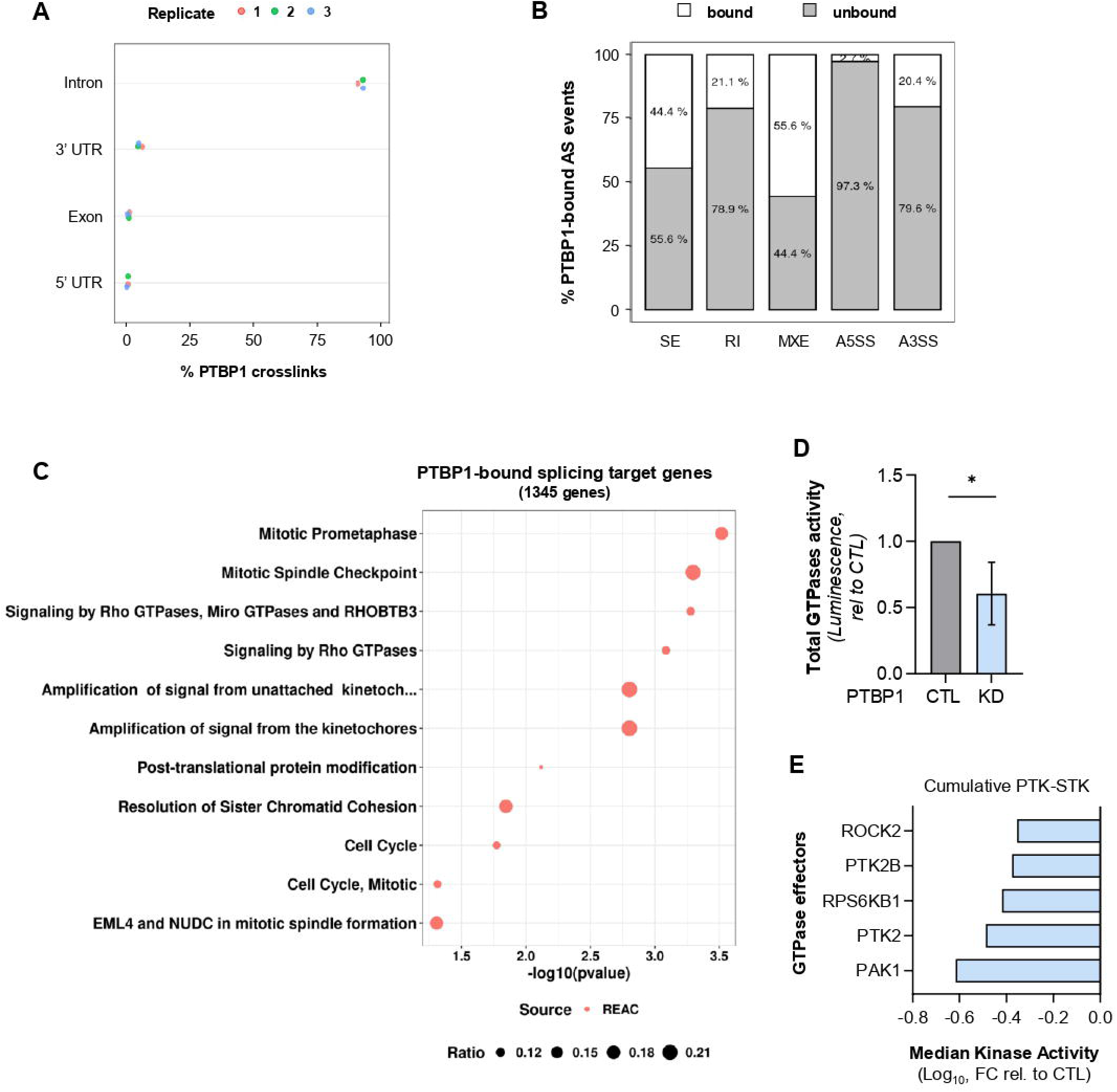
Identification of Rho GTPase signaling as a central pathway directly regulated by PTBP1 in AML. **(A)** Positional distribution of PTBP1 binding sites (crosslinks) across transcript features identified by PTBP1-targeted iCLIP in MOLM-14 AML cells (crosslinks detected in at least 2 out or 3 technical replicates, FDR ≤ 0.05 and Extended Score ≥ 3). **(B)** Proportion of PTBP1-bound *vs.* unbound AS events in MOLM-14 cells, determined by integrating PTBP1-directed iCLIP and RNAseq data (MOLM-14 PTBP1 KD *vs.* CTL). **(C)** Pathway enrichment analysis of predicted direct PTBP1 splicing target genes (N = 1 345), performed using GProfiler with Reactome as the source database. Genes were defined as predicted direct targets if they displayed differential alternative splicing (AS) events in the PTBP1 KD *vs.* CTL condition according to RNA-seq analysis, and if the corresponding AS events were bound by PTBP1 based on iCLIP data. **(D)** GTPase activity assay on PTBP1 KD *vs.* CTL MOLM-14 cells. ShRNA-expressing cells were FACS-sorted at day 3 after transduction and the assay was performed at day 6. Luminescence values are normalized to CTL condition. N=4 **(E)** Kinase activity profiling of selected GTPase effector kinases. Protein lysates from PTBP1 KD and CTL MOLM-14, MV4-11 and U937 AML cell lines were used to measure Serine/Threonine kinase (STK) and tyrosine kinase (TK) activity on PamChip peptide microarrays. Median kinase activity in PTBP1 KD condition relative to CTL (= Median Kinase Statistics) is shown for the indicated Rho GTPase effector kinases. For the three AML cell lines, PTBP1 knockdown (KD) conditions were pooled (N=2 per cell line), and the same pooling was applied to the control (CTL) conditions. N=6.

Overall, 1345 differentially spliced genes were identified as putative direct PTBP1 targets, including known targets such as *PTBP2*(37), *CDC42*(24,38), *PKM*(35,36), and *RTN4*(39). Over-representation analysis revealed significant enrichment for pathways related to cell cycle regulation and Rho GTPase signaling (Fig. 4C). Consistently, PTBP1 silencing resulted in reduced global GTPase activity (Fig. 4D) and diminished activity of multiple downstream Rho GTPase effector kinases (Fig. 4E).

Protein–protein interaction network analysis of AML-associated PTBP1 targets - defined as genes bound by PTBP1 near PTBP1-dependent splicing events and consistently mis-spliced upon PTBP1 KD in all four cell lines - identified CDC42 as a central regulatory hub (Fig. 5A, Fig. S6). CDC42 is a member of the Rho family of small GTPases, which plays a key role in actin cytoskeleton organization, cell polarity, vesicle trafficking, cell cycle progression, and protein translation(40,41).

**Figure 5.**
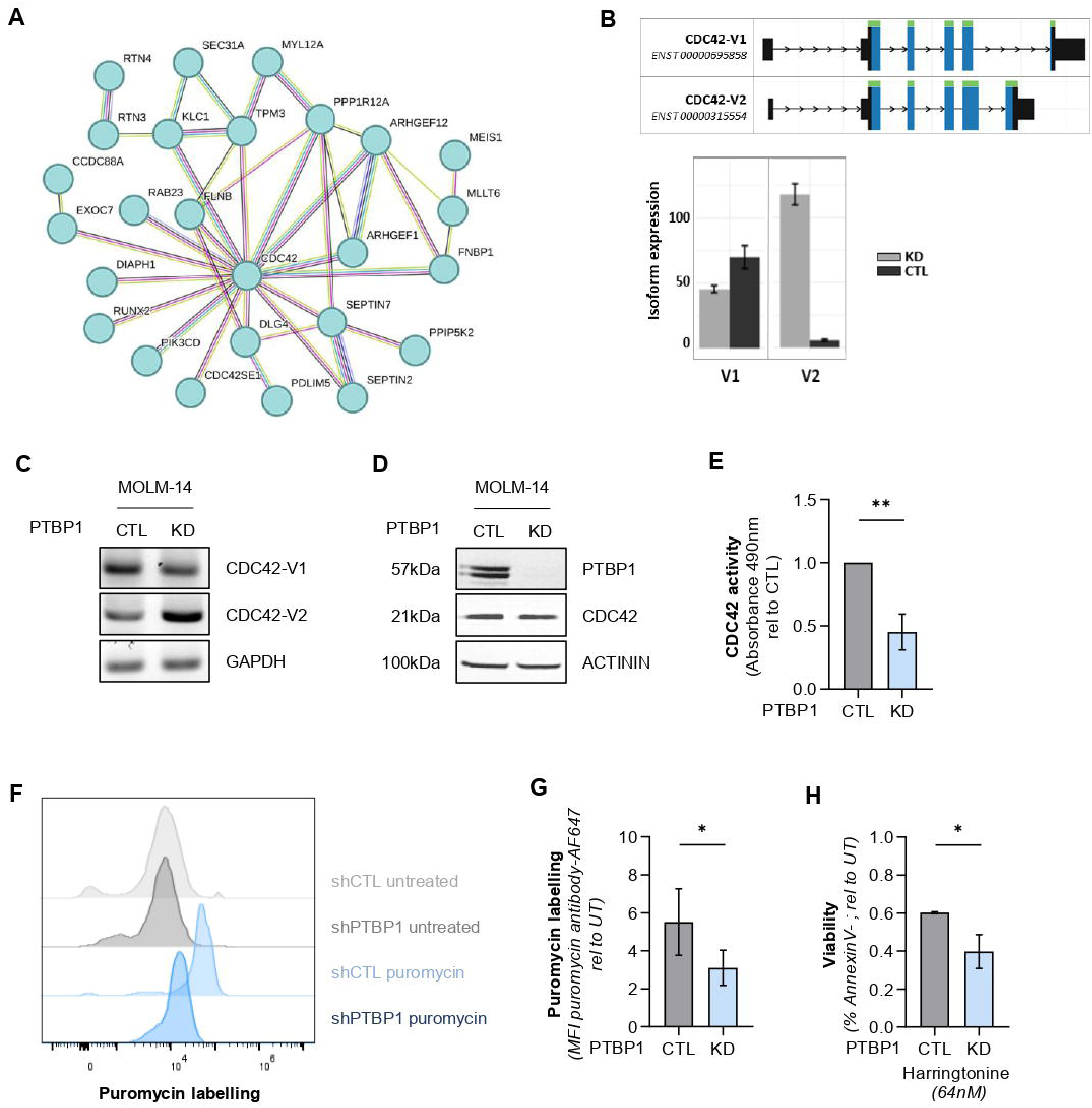
PTBP1 controls CDC42 alternative splicing and function. **(A)** Protein–protein interaction (PPI) network analysis performed using STRING (https://string-db.org/) on conserved, predicted direct PTBP1 splicing targets. Genes were included in the network if they were bound by PTBP1 in MOLM-14 cells and exhibited conserved alternative splicing (AS) regulation across all four AML cell lines in the PTBP1 KD *vs.* CTL condition. Only the central hub, including CDC42, is shown here. The full network is presented in Fig. S6. **(B)** CDC42 transcript splice variants (CDC42-v1, ENST00000695858; CDC42-v2, ENST00000315554) (top). Histogram showing the expression of CDC42 splicing isoforms in control (shCTL, black) and PTBP1 KD (shPTBP1, grey) conditions in AML cells (bottom). Analysis was performed using IsoformSwitchAnalyzeR. **(C)** Agarose gel of representative RT-PCR performed using CDC42 isoform-specific primers in PTBP1 KD *vs.* CTL MOLM-14 cells. The housekeeping gene *GAPDH* was used as a loading control. **(D)** Western Blot analysis of CDC42 and PTBP1 protein expression in FACS-sorted GFP^+^ PTBP1 KD and CTL MOLM-14 cells. The housekeeping protein ACTININ was used as a loading control. **(E)** CDC42-specific GTPase activity assay performed on FACS-sorted GFP^+^ PTBP1 KD and CTL MOLM-14 cells. CDC42 GTPase activity was measured by absorbance at 490nm. N=3. **(F-G)** Flow cytometry analysis of puromycin labeling. Representative flow-cytometry profiles of puromycin incorporation **(F)** and histogram depicting mean puromycin staining fluorescence intensity **(G)** in shPTBP1-*vs.* shCTL-expressing MOLM-14 cells are shown. N=5. **(H)** ShPTBP1- and shCTL-expressing MOLM-14 cells were treated for 24h with the inhibitor of protein translation initiation Harringtonine at 64nM. Viability was assessed by flow cytometric measurement of AnnexinV staining. N=3. GFP^+^ shRNA-expressing cells used for the assays in **(C-H)** were FACS-sorted at day 3 after transduction and all the analyses were performed at day 6.

The activation of Rho GTPases requires their switch from the inactive GDP-bound form to an active GTP-bound form, a process which is promoted by the Rho guanine nucleotide exchange factor (RhoGEF) family. Notably, several RhoGEFs, including ARHGEF1 and ARHGEF12, also exhibited PTBP1-dependent splicing alterations, suggesting multi-level regulation of the Rho GTPase pathway by PTBP1 (Fig. 5A).

In-depth *in silico* analyses, validated by isoform-specific PCR and/or immunoblotting, demonstrated that PTBP1 KD induces a splicing switch in both *ARHGEF1* and *CDC42* in AML cells (Fig. S7A-C, Fig. 5B-D). In particular, PTBP1 depletion resulted in the up-regulation of CDC42-v2 isoform at the expense of CDC42-v1 (Fig. 5B-C), a shift previously reported in other cancers and physiological contexts(38,42). Notably, this isoform switch did not alter total CDC42 protein levels (Fig. 5D), but led to a marked reduction in CDC42 GTPase activity in AML cells (Fig. 5E). Although these two CDC42 splicing isoforms differ only in their C-terminal region, they have been shown to have distinct functional properties(24,38,43,44). Accordingly, PTBP1-loss–driven upregulation of CDC42-v2 was previously shown to inhibit protein synthesis in murine hematopoietic cells(24). In line with this finding, PTBP1-deficient AML cells exhibited reduced global protein synthesis, as measured by puromycin incorporation (Fig. 5F-G), and increased sensitivity to the translation initiation inhibitor harringtonine (Fig. 5H).

Collectively, these results indicate that PTBP1 regulates a Rho GTPase–centered splicing program that sustains AML cell fitness and growth by modulating GTPase signaling and protein synthesis.

### Pharmacological inhibition of CDC42 selectively impairs AML cell survival and enhances venetoclax efficacy

Our results suggest that PTBP1 is a key regulator of AML cell growth through its control of CDC42 splicing and function. Given the challenges of directly targeting splicing factors like PTBP1 in the clinic, we explored the potential of CDC42 as a downstream therapeutic target. Pharmacological inhibition of CDC42 using apigenin (a flavonoid shown to inhibit CDC42 activity by targeting ARHGEF1(45)) or the selective CDC42 inhibitor CASIN significantly reduced CDC42 GTPase activity (Fig. 6A,C) and global protein synthesis (Fig. 6B,D; Fig. S8A-B). Remarkably, CDC42 inhibition selectively induced cell death *in vitro* in both AML cell lines (Fig. 6E) and primary AML samples (Fig. 6F), while largely sparing healthy peripheral blood mononuclear cells (PBMCs) at the same concentrations (Fig. 6G).

**Figure 6.**
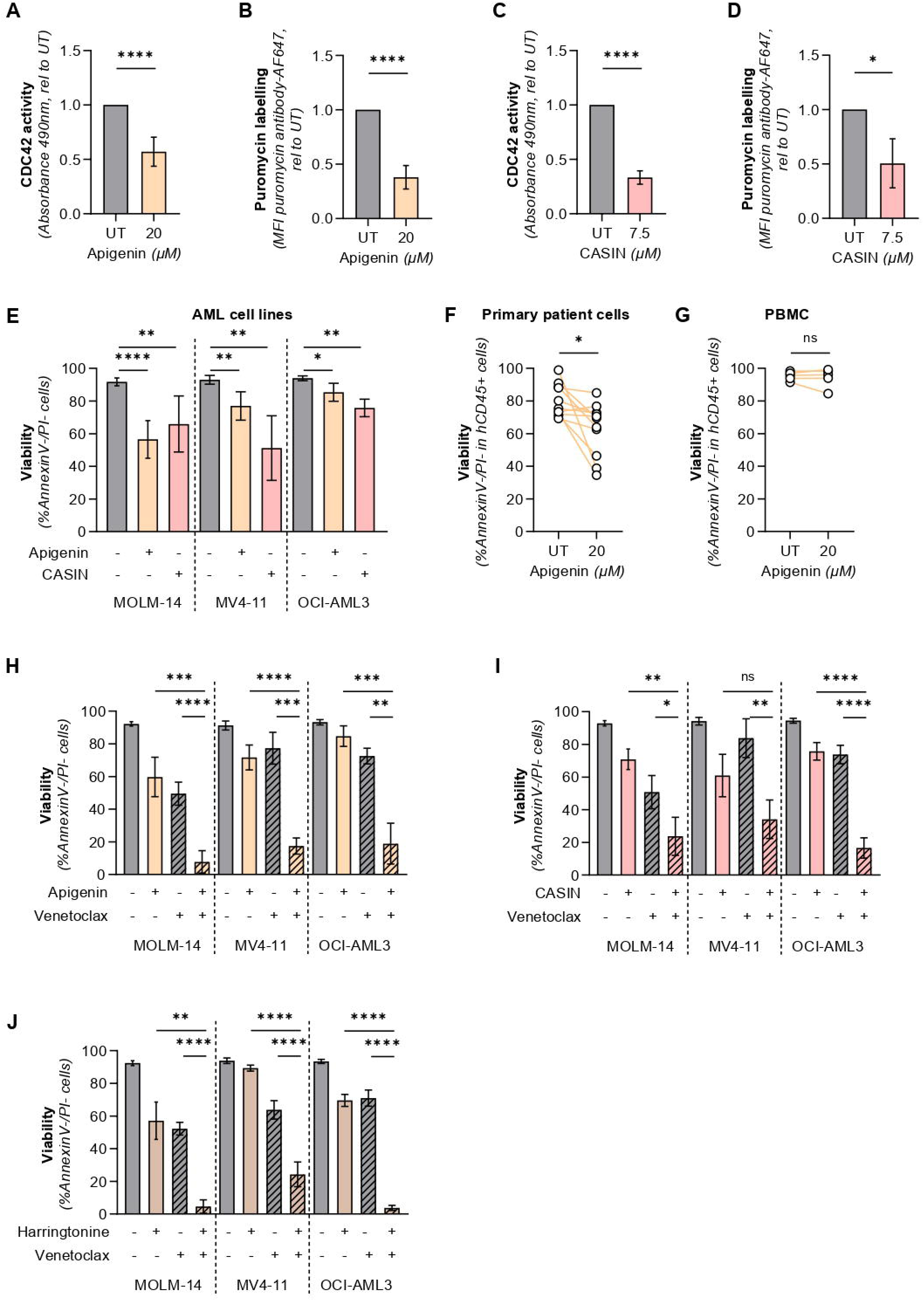
Pharmacological inhibition of CDC42 induces cell death associated with protein synthesis inhibition and potentiates the response to Venetoclax in AML cells. **(A, C)** CDC42-specific GTPase activity assay in MOLM-14 cells treated for 24 h with apigenin (20 μM) **(A)** or CASIN (7.5 μM) **(C)**. CDC42 activity was measured by absorbance at 490 nm. N=5 for apigenin, N= 3 for CASIN. **(B, D)** Flow cytometry analysis of puromycin incorporation in MOLM-14 cells treated for 24 h with apigenin (20 μM) **(B)** or CASIN (7.5 μM) **(D)**. Histograms depict mean puromycin fluorescence intensity for each treatment. N=4 for apigenin, N=3 for CASIN. **(E)** MOLM-14, MV4-11, and OCI-AML3 cells were treated for 24 h with apigenin (20 μM) or CASIN (7.5 μM), as indicated. Viability was assessed by flow cytometric measurement of Annexin V and propidium iodide (PI) staining. N=4-12. **(F)** Primary AML cells (TUH) were treated for 24 h with apigenin (20 μM), and viability was measured by Annexin V/PI staining. **(G)** Peripheral blood mononuclear cells (PBMCs) from healthy donors were treated for 24 h with apigenin (20 μM), and viability was assessed by cytofluorimetric analysis of Annexin V/PI staining. **(H-J)** MOLM-14, MV4-11, and OCI-AML3 cells were treated for 24 h with apigenin (20 μM) **(H)**, CASIN (7.5 μM) **(I)**, or harringtonine (64 nM) **(J)** as single agents or in combination with Venetoclax (0.5 μM), as indicated. N=3-5. Viability was assessed by cytofluorimetric analysis of Annexin V/PI staining.

Venetoclax-based regimens are a standard of care for older or unfit AML patients. Recent research has shown that targeting protein synthesis pathways can sensitize AML cells to venetoclax, mitigating resistance mechanisms(8). Given the role of the PTBP1-CDC42 axis in regulating protein synthesis in AML cells, we thus investigated whether CDC42 inhibition could enhance venetoclax efficacy. Remarkably, co-treatment of AML cells with Venetoclax and either apigenin or CASIN led to a marked increase in Venetoclax-induced cell death *in vitro* (Fig. 6H-I). Consistent with CDC42 playing a key role in modulating translation in AML cells, treatment with the translation inhibitor harringtonine similarly potentiated venetoclax cytotoxicity (Fig. 6J).

Together, these findings identify CDC42 as a therapeutically actionable downstream effector of PTBP1 and support combinatorial strategies integrating CDC42 inhibition with venetoclax to selectively target AML cells.

## DISCUSSION

Acute myeloid leukemia (AML) remains one of the most aggressive hematologic malignancies, characterized by high relapse rates and poor overall survival despite intensive therapy. Aberrant RNA splicing is increasingly recognized as a hallmark of AML and a potential therapeutic vulnerability. However, its clinical exploitation has been limited by challenges such as the risk of toxicity associated with global splicing perturbation, the difficulty of selectively targeting splicing regulators *in vivo*, the lack of patient stratification strategies, and an incomplete understanding of how splicing-directed therapies can synergize with existing treatments(46). To overcome these obstacles, a deeper understanding of AML-specific splicing networks and their critical downstream effectors is essential.

PTBP1, a well-characterized RNA-binding protein and splicing factor, has been linked to tumorigenesis and adverse prognosis across multiple solid malignancies(20). Although its role in hematologic cancers has been less extensively studied, emerging evidence implicates PTBP1 in distinct aspects of AML biology(26–29). Here, we demonstrate that AML cells exhibit a marked dependency on PTBP1 for their growth and survival.

Mechanistically, PTBP1 repression in leukemic cells triggers extensive reprogramming of alternative splicing, affecting thousands of genes.

While PTBP1 has also been reported to regulate mRNA stability and translation through binding to 5′ and 3′ untranslated regions(20,47–49) and, more recently, transcription via co-localization with the master hematopoietic transcription factor RUNX1(29), we observed only modest and heterogeneous changes in transcript abundance following PTBP1 knock-down. Supported by iCLIP data demonstrating predominant intronic binding, our findings suggest that PTBP1 supports AML maintenance primarily through its role as a splicing regulator, rather than through global control of mRNA abundance or transcription.

Among the limited number of genes consistently altered at the expression level upon PTBP1 depletion, PTBP1 paralog PTBP2 emerged as a notable target. Consistent with the well-described mechanism by which PTBP1 represses PTBP2 via alternative splicing-coupled nonsense-mediated decay(50), PTBP1 loss resulted in robust PTBP2 upregulation at both the transcript and protein levels. However, despite reports that PTBP2 can partially compensate for PTBP1 loss in certain contexts(25,37,51), its induction was insufficient to rescue the essential functions of PTBP1 in AML cells.

Previous studies have linked PTBP1 to metabolic regulation in AML(28,29), particularly through control of PKM alternative splicing and transcriptional regulation of key glycolytic genes. While we observed the expected PKM splicing switch upon PTBP1 repression, downstream effects on glycolysis and oxidative phosphorylation were heterogeneous across AML models and did not correlate with growth inhibition, suggesting that altered glucose metabolism is unlikely to represent the dominant mechanism underlying PTBP1 dependency in AML.

On the other hand, integrated analysis of PTBP1-dependent splicing changes and iCLIP data uncovered an AML-associated PTBP1 splicing signature enriched for genes involved in cell division, endocytosis, and vesicle trafficking. Notably, PTBP1 direct targets were strongly enriched for regulators of Rho GTPase signaling, with CDC42 emerging as a central node within this network. The small GTPase CDC42 plays critical roles in the control of cell polarity, cytoskeletal dynamics, vesicle trafficking, and proliferation(52–56), and its dysregulation has been linked to cancer(57). Further complexity arises from alternative splicing of CDC42, generating two isoforms (CDC42-v1 and CDC42-v2) with distinct functional properties(24,38,43). While PTBP1-mediated repression of the CDC42-v2 isoform has been described in other cellular systems(24), its relevance in leukemia has not been explored.

Here, we show that PTBP1 depletion induces a conserved shift toward CDC42-v2 expression in AML cells, accompanied by impaired protein synthesis and heightened sensitivity to translational inhibition. These findings suggest that PTBP1 sustains leukemia cell growth, at least in part, by modulating protein translation through CDC42 isoform regulation. Consistent with this model, similar translational defects have been reported upon PTBP1 loss or CDC42-v2 overexpression in normal murine hematopoietic cells^20^, indicating that this regulatory axis may operate in both normal and malignant hematopoiesis.

Beyond translational control, precise regulation of CDC42 activity is essential for hematopoietic stem cell (HSC) polarity and function, and its dysregulation has been linked to aging and leukemogenesis(58–60). In this context, elevated CDC42 function promotes leukemia-initiating cell fate at the expense of differentiation(60). Our data show that PTBP1 knockdown impairs clonogenicity and leukemogenic potential *in vivo*, raising the possibility that PTBP1-mediated CDC42 regulation contributes to leukemia stem cell function and maintenance.

Importantly, our findings establish that AML cells are functionally dependent on both PTBP1 and CDC42 activity. In line with this, pharmacological inhibition of CDC42 selectively induced cell death in AML cell lines and primary patient samples while largely sparing normal hematopoietic cells *in vitro*. Given the central role of Rho GTPases in cancer, these findings identify the CDC42 as a tractable therapeutic vulnerability in AML.

Current AML treatment relies on intensive cytarabine–anthracycline chemotherapy for fit patients and venetoclax-based regimens for older or unfit individuals, a population characterized by particularly poor outcomes(2–4). Recent work has highlighted the importance of RNA splicing in modulating AML sensitivity to the BCL-2 inhibitor venetoclax(6,7). In parallel, venetoclax exposure has been shown to trigger an adaptive increase in protein synthesis in AML cells, and disrupting this adaptive response by perturbing translation enhances treatment sensitivity(8). In agreement with these observations, we demonstrate that combining venetoclax with either the protein synthesis inhibitor harringtonine or CDC42 inhibitors dramatically increases AML cell sensitivity to the treatment, suggesting a rational combination strategy that exploits AML dependency on the PTBP1–CDC42 axis.

**Collectively, our findings identify PTBP1 as a broad and non-redundant molecular dependency in AML, where it acts as a critical regulator of CDC42 signaling and protein translation. Furthermore, our work proposes a new rational therapeutic combination that could be leveraged to enhance the efficacy of venetoclax-based regimens.**

## Supporting information

Supplementary Information, including Supplementary Figures and their legends, and Supplementary Materials and Methods.

## ACKNOWLEDGEMENTS

We thank CREFRE for mice experimentation. We are grateful to Manon Farcé for her assistance with flow cytometry, and to Estelle Saland and Constance Manso for their support with *in vivo* experimentation. We thank Anne-Marie Bénot, Jérôme Lacan and Latifa Jarrou for their administrative support. Additionally, we extend our gratitude to Johannes Zuber and Johannes Schmoellerl (Research Institute of Molecular Pathology, IMP, Vienna, Austria) for kindly providing the viral constructs used in this study and for their valuable scientific discussions.

This work was supported by the INCa (PLBIO22-032), the Ligue Nationale de Lutte contre le Cancer, the ARC, and a career development grant by Gilead Sciences.

## AUTHORS CONTRIBUTIONS

MO, ML and MG conceived of and designed the experiments. MO and ML performed the experiments, with assistance from MF, AGr, AB and MG for specific procedures and analyses. SB, FV, CR, VDM provided patients’ samples. YA performed most of the bioinformatics analyses, with assistance from AGa. AS assisted with mice experimentation. CL, JES, LP and CJ made intellectual contributions. MDM made intellectual contribution and designed and performed iCLIP analyses. VP made intellectual contribution and provided expert guidance for the bioinformatics analysis. MG and MO wrote the manuscript, with significant contributions of ML for figures preparation.

## DATA AVAILABILITY STATEMENT

The RNA sequencing and iCLIP datasets generated during the current study have been deposited in the Gene Expression Omnibus (GEO) and will be made publicly available upon publication of the manuscript. The datasets are available from the corresponding author upon reasonable request.

