## Supplementary Information, including Supplementary Figures and their legends, and Supplementary Materials and Methods. for "A PTBP1–CDC42 splicing axis regulates leukemia growth and venetoclax sensitivity in acute myeloid leukemia"

#### Figure S1

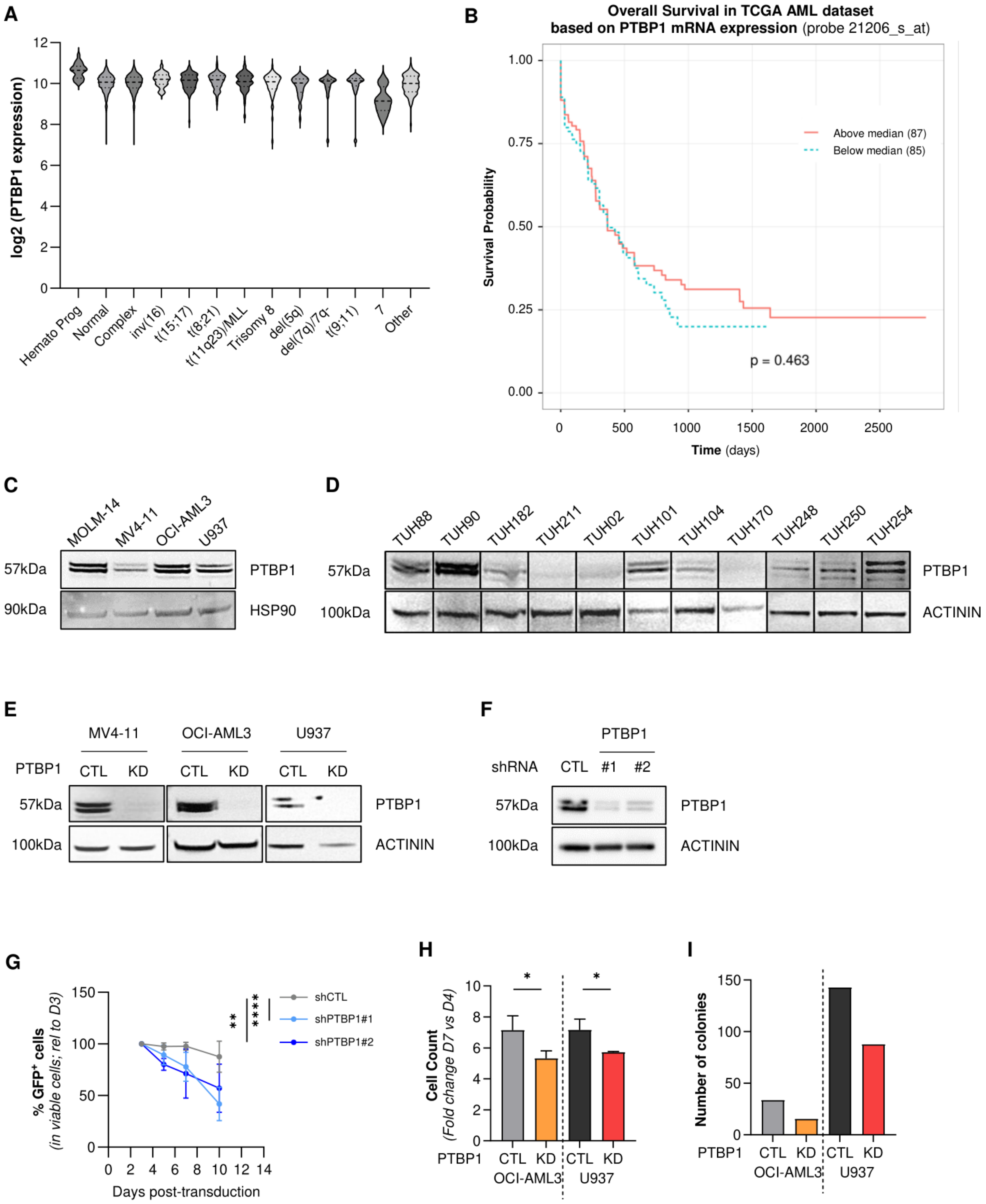

**Figure S1. (A)** PTBP1 transcript expression in AML patients stratified by karyotype. Gene expression data pooled by BloodSpot ("BloodSpot: a database of gene expression profiles and transcriptional programs for healthy and malignant haematopoiesis". Nucl. Acids Res. 2015) were obtained from GSE13159, GSE15434, GSE61804, GSE14468, and The Cancer Genome Atlas (TCGA). **(B)** Overall survival of AML patients stratified by median PTBP1 expression (red line, above median; dotted blue line, below median). Gene expression and clinical data were obtained from TCGA via BloodSpot. **(C, D)** Western blot analysis of PTBP1 protein expression in human AML cell lines (MOLM-14, MV4-11, OCI-AML3, U937) **(C)** and primary AML patient samples **(D)**. HSP90 in **(C)** and ACTININ in **(D)** were used as a loading controls. **(E-F)** Western blot analysis of PTBP1 expression in GFP-positive, FACS-sorted MV4-11, OCI-AML3 and U937 **(E)** or MOLM-14 cells **(F)** 6 days after transduction with SGEN lentiviral vectors expressing shRNAs targeting PTBP1 (shPTBP1#1 or shPTBP1#2) or a control shRNA (shCTL), as indicated. ACTININ was used as a loading control. ShPTBP1#1 was used in the experiments shown in Fig. S1E and Fig.1-4. **(G)** Competitive proliferation assay on MOLM-14 cells transduced with lentiviral vectors expressing shPTBP1#1, shPTBP1#2 or shCTL. A mixed AML population containing GFP-positive (shRNA-expressing) and GFP-negative (untransduced) cells was analyzed. The percentage of viable GFP-positive cells was monitored over time by flow cytometry and normalized to day 0. N=7 **(H)** Trypan blue staining-based cell counting of PTBP1 KD *vs.* CTL AML cells. Viable shRNA-expressing ($PI^-/GFP^+$) AML cells were FACS sorted 3 days after transduction and plated at the same concentration. Viable cell counts at day 7 were normalized by input counts at day 4. N=3. **(I)** Methylcellulose colony forming assay on GFP-positive (shRNA-expressing) AML cells. FACS-sorted GFP-positive cells were plated in semi-solid medium 3 days after transduction. The number of colonies was assessed by MTT staining 10 days after plating. N=1. Data are presented as mean values ± SD of biological replicates.

#### Figure S2

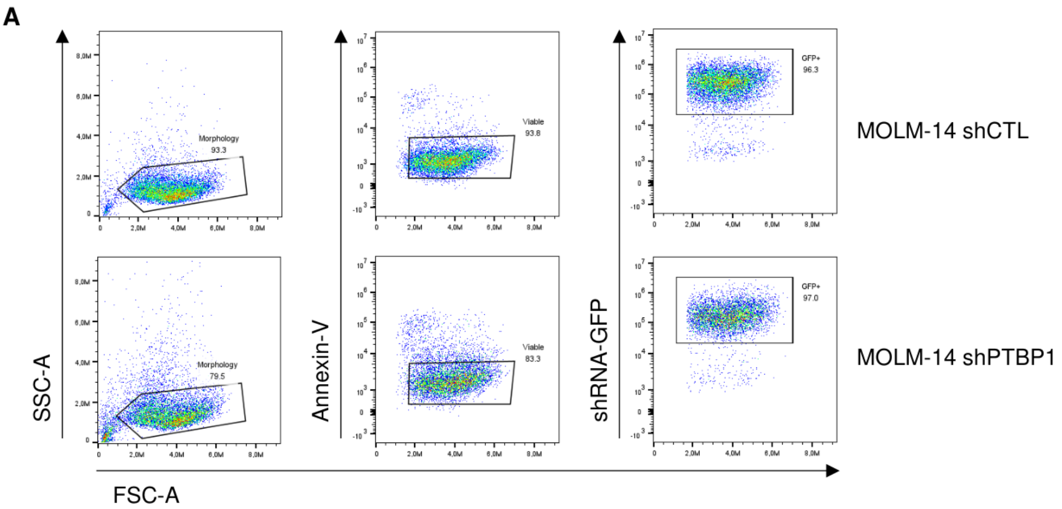

**Figure S2. (A)** AML cells were transduced with SGEN lentiviral vectors expressing either a control shRNA (CTL) or a PTBP1-targeting shRNA (KD), both coupled to GFP. GFP-positive cells were sorted at day 3 after transduction to a purity >95% and injected intravenously on the same day into immunodeficient NOD/SCID mice (Fig.1A). Assessment of the percentage of GFP-positive MOLM-14 cells after FACS sorting prior to injection into recipient animals is shown.

### Figure S3

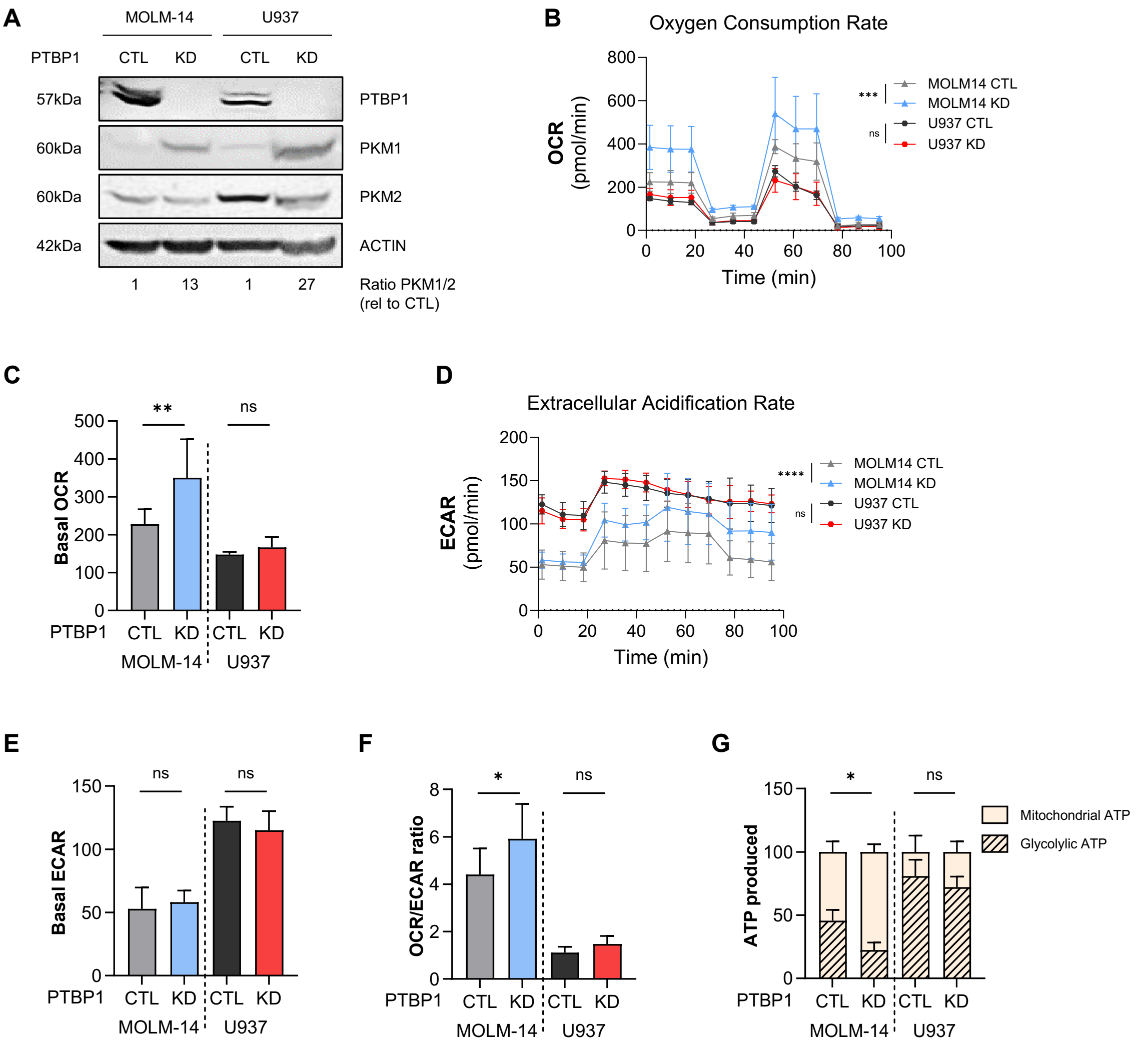

**Figure S3.** MOLM-14 and U937 cell lines were transduced with miRE-based SGEN lentiviral vectors expressing either a control shRNA (CTL) or a PTBP1-targeting shRNA (KD), both coupled to GFP. GFP-positive cells were sorted at day 2 after transduction and analysis of PKM expression and metabolic parameters was performed at day 6. **(A)** Western blot analysis of protein expression was performed in PTBP1-knockdown (KD) and control (CTL) MOLM-14 and U937 AML cells to assess PTBP1 as well as the PKM1 and PKM2 splice isoforms. The PKM1/PKM2 ratio, determined by densitometric analysis of the bands, is indicated. β-ACTIN was used as a loading control. **(B, D)** Representative Seahorse XF Mitochondrial Stress Test profiles showing oxygen consumption rate (OCR), reflecting mitochondrial respiration and oxidative phosphorylation activity **(B)**, and extracellular acidification rate (ECAR), reflecting glycolysis and lactate production, **(D)** in GFP-positive AML cells expressing PTBP1-targeting shRNAs or a negative control, as indicated. **(C, E, F)** Quantification from Seahorse Mitochondrial Stress Tests showing basal OCR **(C)**, basal ECAR **(E)**, and the OCR/ECAR ratio **(F)**. N=4 for MOLM-14, N=3 for U937. **(G)** ATP quantification assay determining the proportion of ATP derived from mitochondrial oxidative phosphorylation *vs.* glycolysis in PTBP1 KD *vs.* CTL MOLM-14 and U937 cells. N=3 for MOLM-14, N=3 for U937.

### Figure S4

A

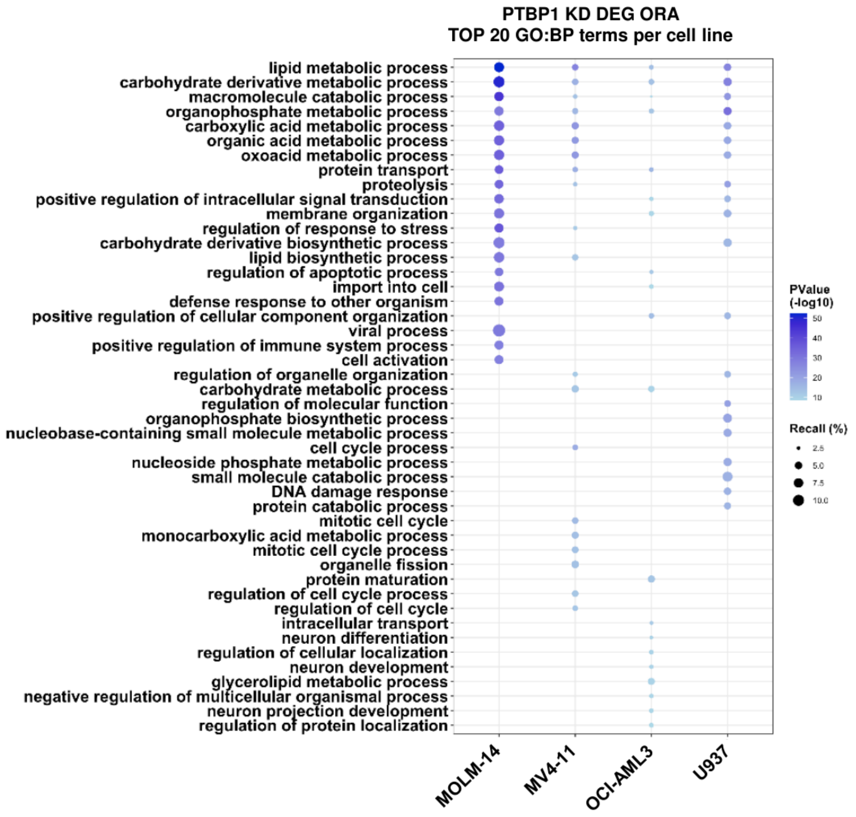

B

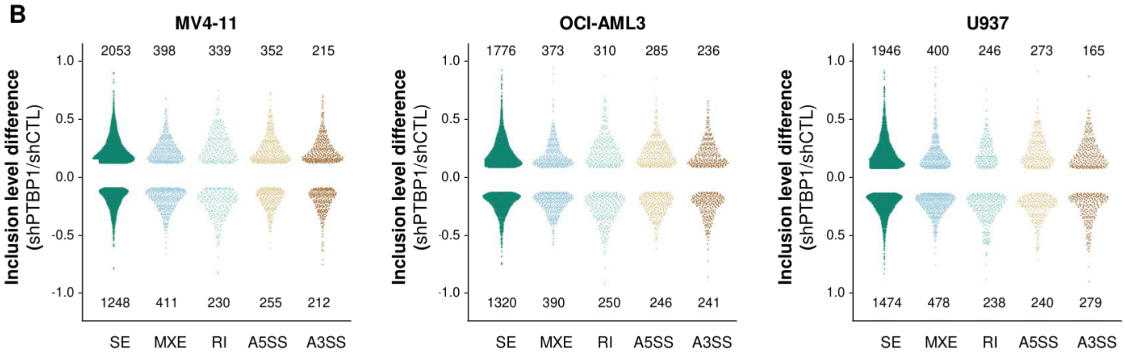

C

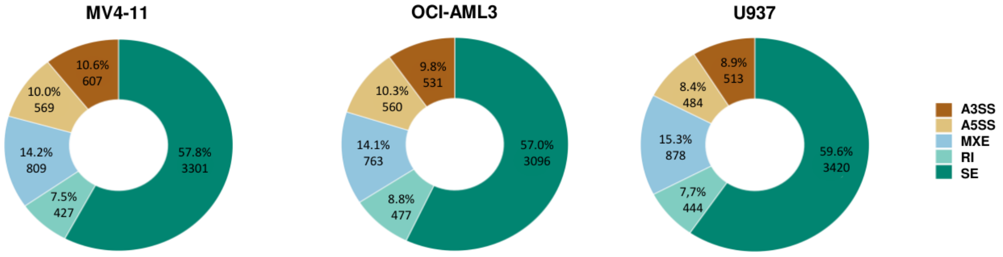

#### Figure S4

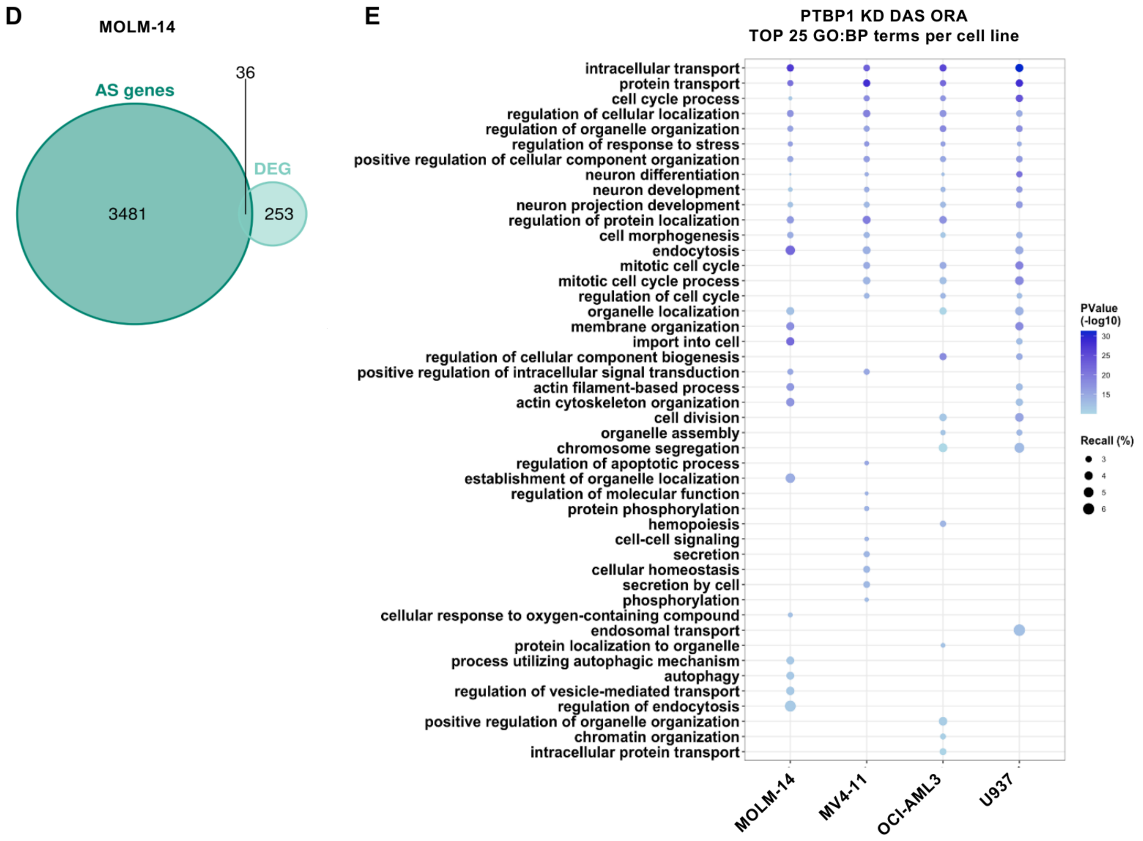

**Figure S4. (A)** Top 20 enriched pathways independently identified from comparative over-representation analysis (ORA) of the lists of differentially expressed genes (DEGs) in the PTBP1-knockdown (KD) *vs.* control (CTL) condition in MOLM-14, MV4-11, OCI-AML3, and U937 cells (FDR ≤ 0.05; MOLM-14: 1191 genes, MV4-11: 558 genes, OCI-AML3: 484 genes, U937: 754 genes). Gene Ontology Biological Processes (GO:BP) was used as the source database for pathway enrichment analysis. **(B)** Beanplot representing the distribution of alternative splicing (AS) events inclusion level difference in PTBP1 KD *vs.* CTL AML cells, binned by AS event type (SE: Skipped Exon, MXE: Mutually Exclusive Exon, RI: Retained Intron, A5SS and A3SS : Alternative 5’ or 3’ Splice Site). AS events with Incl. Level. Diff ≥ 0.1, FDR ≤ 0.05, avg. read counts>10 are shown and numbers are indicated on the plot. Differential AS events were identified by rMATS and filtered using maser R package. Data relative to MV4-11, OCI-AML3 and U937 cell lines are shown. **(C)** Donut charts illustrating the distribution of differential AS events across specific AS event types in PTBP1 KD *vs.* CTL MV4-11, OCI-AML3 and U937 cells, as indicated. Both the absolute number and relative proportion of each AS event type are shown. **(D)** Venn diagram representing the overlap between differentially AS and differentially expressed genes identified in PTBP1 KD *vs.* CTL MOLM-14 cells. **(E)** Top 25 enriched pathways independently identified from comparative over-representation analysis (ORA) of the lists of differentially AS genes (top 500 events, all AS types) in the PTBP1-knockdown (KD) *vs.* control (CTL) condition in MOLM-14, MV4-11, OCI-AML3, and U937 cells. Gene Ontology Biological Processes (GO:BP) was used as the source database for pathway enrichment analysis.

#### Figure S5

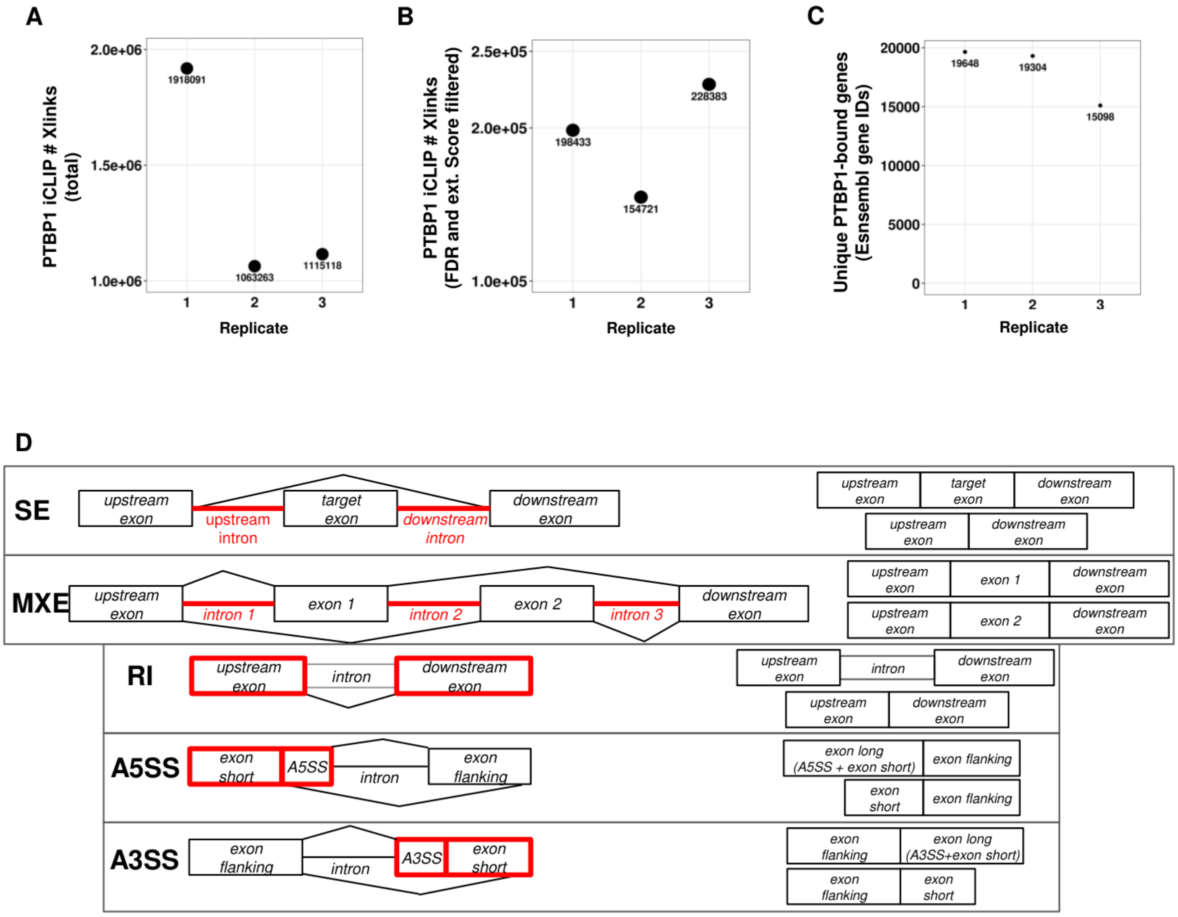

**Figure S5. (A)** Total number of PTBP1 cross-links (Xlinks) identified on the whole transcriptome. **(B)** Total number of PTBP1 cross-links (Xlinks) filtred by FDR ≤ 0.05 and extended score ≥3. **(C)** Total number of PTBP1 Cross-links (Xlinks) associated with an Ensembl gene ID. **(D)** Schematic representation of the strategy used to identify AS events directly bound by PTBP1. Elements in red represent the genomic coordinates of AS events in which overlap with PTBP1 iCLIP crosslink sites was assessed to infer direct binding and regulation.

#### Figure S6

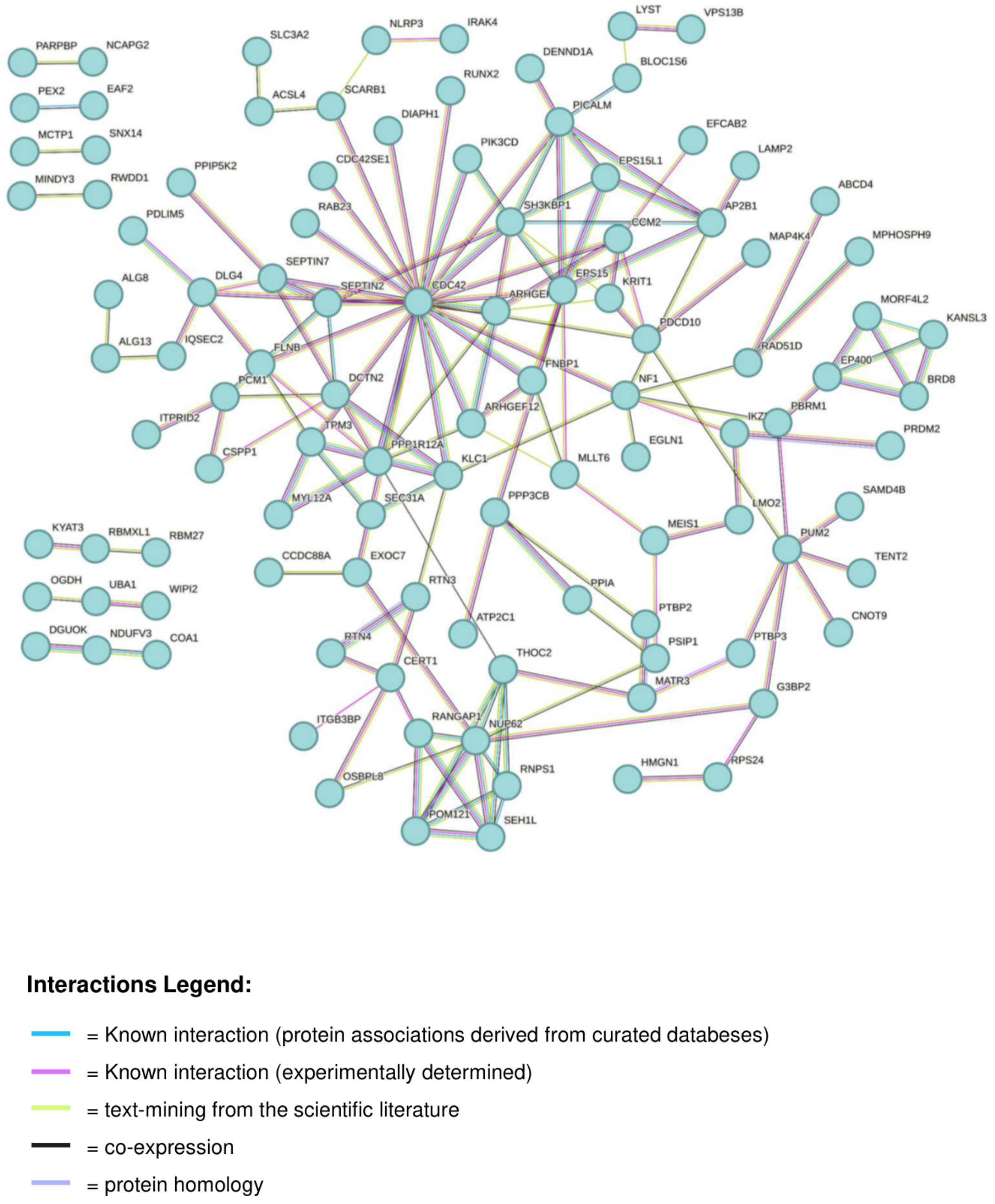

**Figure S6.** Protein–protein interaction (PPI) network analysis performed using STRING (https://string-db.org/) on conserved, predicted direct PTBP1 splicing targets (186 genes). Genes were included in the network if they were bound by PTBP1 in MOLM-14 cells and exhibited conserved alternative splicing (AS) regulation across all four AML cell lines in the PTBP1 KD *vs.* CTL condition. Disconnected nodes in the network were hidden.

#### Figure S7

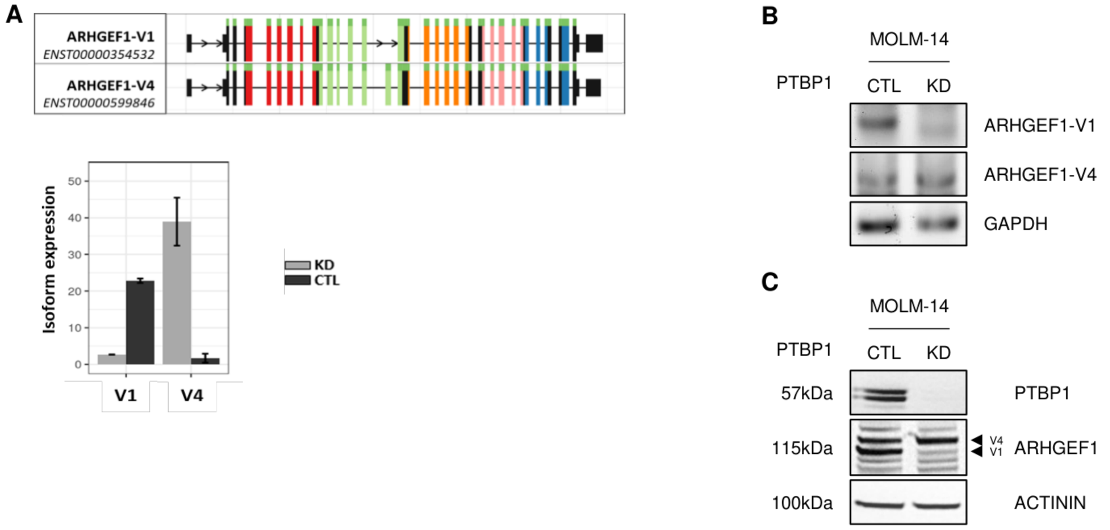

**Figure S7. (A)** ARHGEF1 transcript splice variants (ARHGEF1-v1, ENST00000354532; ARHGEF1-v4, ENST00000599846) (top). Histogram showing the expression of ARHGEF1 splicing isoforms in control (shCTL, black) and PTBP1 KD (shPTBP1, grey) conditions in AML cells (bottom). Analysis was performed using IsoformSwitchAnalyzeR. **(B)** Agarose gel of representative RT-PCR performed using ARHGEF1 isoform-specific primers in PTBP1 KD *vs.* CTL MOLM-14 cells. The housekeeping gene *GAPDH* was used as a loading control. **(C)** Western Blot analysis of ARHGEF1 and PTBP1 protein expression in FACS-sorted GFP$^{+}$ PTBP1 KD and CTL MOLM-14 cells. The housekeeping protein ACTININ was used as a loading control.

#### Figure S8

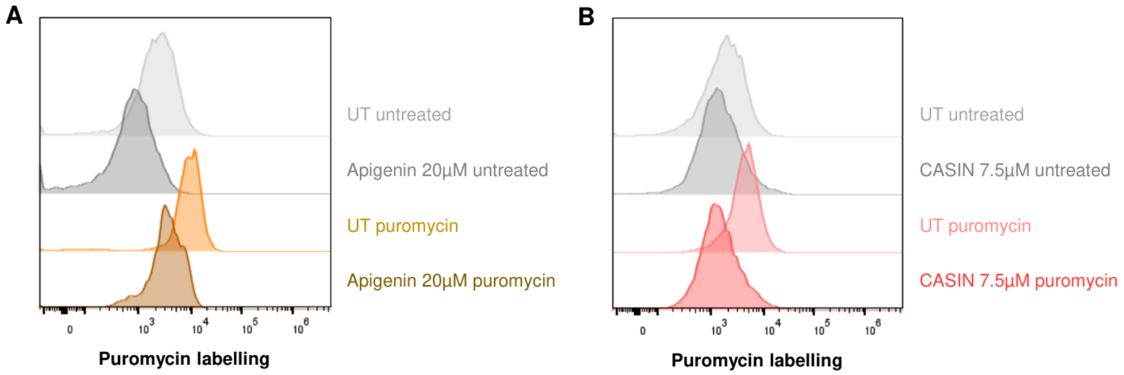

**Figure S8. (A, B)** Representative flow-cytometry profiles of puromycin incorporation in MOLM-14 cells treated for 24 h with apigenin (20 µM) **(A)** or CASIN (7.5 µM) **(B).**

#### SUPPLEMENTARY MATERIAL

##### AML cell lines

All AML cell lines were mantained in Minimum Essential Medium (MEM)-α (Gibco, Thermofisher Scientific), supplemented with 5% heat-inactivated fetal bovine serum (FBS, Invitrogen), 100 U/mL penicillin, and 100µg/mL streptomycin (Life Technology). Cells were passaged every 2–3 days to maintain exponential growth and were routinely tested for mycoplasma contamination.

Human epithelial kidney (HEK) 293T/17 (ATCC) were cultured in Dulbecco's Modified Eagle Medium (DMEM) GlutaMAX™ (Gibco, Thermofisher Scientific), supplemented with 10% FBS and 100 U/mL penicillin, and 100µg/mL streptomycin (P/S). All cells were incubated at 37°C in a 5% $CO_2$ atmosphere.

##### Primary AML and hematopoietic cells

Primary AML cells were obtained from peripheral blood samples collected during routine diagnostic procedures at Toulouse University Hospital (TUH) after obtaining informed consent from patients. If not used immediately, cells were stored at the Hémopathies Inserm Midi-Pyrénées (HIMIP) collection (BB-0033-00060). In accordance with French legislation, the HIMIP collection has been declared to the Ministry of Higher Education and Research (DC 2008-307 collection 1) and a transfer agreement (AC 2008-129) has been obtained following approval by the Comité de Protection des Personnes Sud-Ouest et Outremer II (ethical committee).

Primary AML cells and peripheral blood mononuclear cells (PBMCs) were isolated by Ficoll-Paque density gradient centrifugation (GE Healthcare). Primary cells were cultured at 37 °C with 5% $CO_2$ in Iscove's Modified Dulbecco's Medium with GlutaMAX (IMDM, Gibco, Life Technologies), supplemented with 20% fetal calf serum and 100 U/mL penicillin and 100 µg/mL streptomycin (Life Technology).

**Measurement of Oxygen Consumption and Extracellular Acidification Rate using Seahorse Assay**

Oxygen Consumption Rate (OCR) and Extra-Cellular Acidification Rate (ECAR) were measured using the Seahorse XFe24 Analyzer (Agilent Technologies) and the Seahorse XF Cell Mito Stress Test. The day before the assay, the sensor cartridge was hydrated in Seahorse XF Calibration Buffer and incubated at 37°C without $CO_2$. Seahorse XFe24 microplate wells were coated with Cell-Tak (Corning, Cat#354240) at a concentration of 22.4µg/mL and incubated overnight at 4°C. AML cells were seeded at a density of $2x10^5$ cells per well in Seahorse XF base Medium supplemented with 5.6mM glucose, 1mM pyruvate and 2mM glutamine. Cells were centrifuged at 80 x g for 5minutes to promote adherence and then incubated for 1 hour at 37°C in a non-$CO_2$ incubator. Initially, baseline cellular OCR was measured, from which basal respiration can be derived by subtracting non-mitochondrial respiration. Next oligomycin, a complex V inhibitor, was added and the resulting OCR was used to derive ATP-linked respiration (by subtracting the oligomycin rate from baseline cellular OCR) and proton leak respiration (by subtracting non-mitochondrial respiration from the oligomycin rate). Next FCCP, a protonophore, was added to collapse the inner membrane gradient, allowing ETC to function at its maximal rate and maximal respiratory capacity was derived by subtracting non-mitochondrial respiration from the FCCP rate. Last, antimycin A and rotenone, inhibitors of complex III and I, were added to shut down ETC function, revealing the non-mitochondrial respiration.

Basal OCR (indicator of baseline mitochondrial respiration) and ECAR (indicator of baseline glycolytic activity) were measured using the Seahorde XFe24 analyzer according to the manufacturer's protocol.

**ATP assay**

Mitochondrial and glycolytic ATP was measured using the CellTiter-Glo Luminescent Cell viability Assay (Promega, Cat#G7570), following manufacturer instructions.

Cells were plated at a concentration of 0.3 x $10^6$ cells/mL, 80 µl/well, in white 96-wells in pre-warmed MEM-alpha medium. Cells were treated with 20 µl/well of either Phosphate Buffer Saline (PBS), sodium iodoacetate (Sigma-Aldrich, Cat.#I2512), oligomycin (Sigma-Aldrich, Cat.#75351) plus antimycin A (Sigma-Aldrich, Cat.#A8674), or oligomycin plus antimycin A plus sodium iodoacetate. After 1 hour of incubation à 37°C, 100µL of CellTiter-Glo reagent was added directly to each well for a final volume of 200µL. Plates were incubated for 10min at room temperature in the dark. Luminescence was recorded using the microplate reader CLARIOstar Plus (BMG Labtech). Luminescence values were normalized to the vehicle. By comparing the different conditions, total ATP and the percentages of mitochondrial and glycolytic ATP were calculated as follows:

<u>Mitochondrial ATP</u>= Luminescence (sodium iodoacetate) – Luminescence (oligomycin + antimycinA + sodium iodoacetate)

<u>Glycolytic ATP</u>= Luminescence (oligomycin + antimycinA) – Luminescence (oligomycin + antimycinA + sodium iodoacetate)

<u>Total ATP</u>= Luminescence (PBS) – Luminescence (oligomycin + antimycinA + sodium iodoacetate)

<u>% Mitochondrial ATP</u>= (Mitochondrial ATP/ Total ATP) * 100

<u>% Glycolytic ATP</u>= (Mitochondrial ATP/ Total ATP) * 100

**Pan-kinase Activity Assay**

Kinase activity was measured using PamGene technology and the PamStation12 system, which enables real-time monitoring of kinase activity through the PamChip® peptide microarray platform. Briefly, cells were lysed in ice-cold M-PER™ Mammalian Protein Extraction Buffer

(Thermo Fisher Scientific, Cat.#78501), supplemented with Halt™ Phosphatase Inhibitor Cocktail (100X, Thermo Fisher Scientific, Cat.#78420) and Halt™ Protease Inhibitor Cocktail, EDTA-free (100X, Thermo Fisher Scientific, Cat.#87785). After 15 mins of incubation on ice, lysates were centrifuged at 16 000g for 15min at 4°C, snap-frozen in liquid nitrogen and stored at -80°C. Protein concentration was determined using BCA Protein Assay (ThermoFisher Scientific, Cat#23225), according to the manufacturer's instructions. Following a step of PamChip pre-processing (performed following the manufacturer's protocol), 2-4µg of protein were mixed with the PTK or STK Basic Mix and loaded on the array. The PamChip peptide microarray was imaged on PamStation12 CCD camera. Spot signal intensities were quantified and corrected for local background using BioNavigator software v6.3 (PamGene International). Upstream Kinase Analysis, a functional scoring method (PamGene) was used to rank kinases based on combined specificity scores (based on peptides linked to a kinase, derived from 6 databases) and sensitivity scores (based on treatment-control differences). Only kinases with a Median Final Activity Score > 1.2 were retained for downstream analysis. The median kinase statistic represents the median fold-change ($Log_{10}$) in kinase activity compared to the control, indicating significant changes in activity between shPTBP1 and shCTL conditions across all three cell lines. Gene names for the selected kinases were identified using the Uniprot database.

**GTPase Activity Assays**

Total GTPase activity was measured using the GTPase-Glo™ Assay (Promega, Cat#V7681), following the manufacturer's instructions with slight modifications. Briefly, cell lysis was performed as described above (see Pan-Kinase Activity Assay protocol) using 100µL of buffer per $1\times10^6$ cells. Luminescent signal corresponding to the amount of GTP remaining after hydrolysis was measured according to the manufacturer's protocols using the plate reader CLARIOstar Plus, BMG Labtech.

CDC42 activity was quantified using the Cdc42 G-LISA GTPase Activation Assay (Cytoskeleton Inc., Cat#BK127), following the manufacturer's instructions. Briefly, cells were lysed in the buffer provided in the kit and clarified by centrifugation. Activated GTP-bound CDC42 was detected via colorimetric ELISA-based method and absorbance measurements was assessed at 490nm.

**RT-PCR Analysis**

Total RNA was extracted using RNeasy Mini kit (Qiagen, Cat#74106) according to the manufacturer's instructions. RNA was used immediately or stored at −80 °C. Reverse transcription into cDNA was performed using the iScript™ Reverse Transcription Supermix (Bio-Rad, Cat#1708891). The expression level of *CDC42* and *ARHGEF1* transcripts was determined by PCR using EconoTaq Plus 2X Master Mix (Lucigen, Cat#30035-1). Variant-specific primers (provided in Supplemental Information) were used to distinguish the splicing isoforms of interest. PCR amplicons were separated by electrophoresis on 1.2% agarose gel prepared in TAE buffer and stained with SYBR™ Safe DNA Gel Stain (Invitrogen, Cat# S33102). DNA bands were visualized using a UV transilluminator on a BioRad ChemiDoc imaging system. Primers sequences are provided in the dedicated table.

**Western Blot Analysis**

Proteins were separated by using 4–12% gradient polyacrylamide SDS–PAGE gels (Life Technologies), and electrotransferred to 0.2 µm nitrocellulose membranes (GE Healthcare). Following incubation with primary antibodies and horseradish peroxidase-conjugated secondary antibodies, immunoreactive bands were detected using enhanced chemiluminescence. Signals were captured with a Pxi camera (Syngene) and analyzed using GeneSys software. Densitometric quantification of immunoblots was performed using GeneTools software. Details on the antibodies used are provided in the dedicated table.

**RNA-seq Differential Expression Analysis**

Raw sequencing files were processed using the nf-core/RNAseq pipeline (v3.10.1) on the Genotoul SLURM HPC cluster. Reads were aligned to the ENSEMBL human genome GRCh38 v108 using STAR (v2.7.9) and quantified with Salmon (v1.9.0)**.** Quality control was performed with FastQC (v0.11.9) and reads were trimmed using Trim Galore (v0.6.7) and Cutadapt (v3.4). Differential gene expression analysis was conducted using DESeq2 (v1.44.0) in R (v4.4.0), with CombatSeq used to adjust for batch effects. Significant changes in gene expression were defined as those with a Benjamini-Hochberg adjusted p-value (padj) ≤ 0.05**.**

For transcriptomics analyses, raw sequencing files were processed using nextflow nf-core/RNAseq pipeline (v3.10.1) [1-2] on Genotoul SLURM high-performance computing (HPC) cluster starting from a samplesheet created using nfcore-rnaseq's "fastq_dir_to_samplesheet.py" script. nfcore/rnaseq pipeline was ran with options -profile genotoul, --igenomes_ignore TRUE, --aligner star_salmon using ENSEMBL human genome GRCh38 v108 sequence (Homo_sapiens.GRCh38.dna.primary_assembly.fa) and annotation (Homo_sapiens.GRCh38.108.gtf) as reference genome options. Briefly, the quality of untrimmed reads was assessed with FastQC (v0.11.9) [3], reads were trimmed with Trim Galore (v0.6.7) [4] and Cutadapt (v3.4) [5]. The quality of the trimmed reads was assessed with FastQC (v0.11.9) using default parameters. Trimmed reads were aligned to the human genome with STAR (v2.7.9) [6] and quantified with Salmon (v1.9.0) [7]. Alignment files were sorted and indexed using SAMtools (v1.16.1) [8], duplicates were marked with PICARD (v2.27.4-SNAPSHOT) [9]. Differential gene expression analysis was performed using R (v4.4.0) and RStudio (build 365) on a x86_64-pc-linux-gnu platform and DESeq2 (v1.44.0) [10]. Salmon quantification files at the transcript level (i.e. quant.sf files) were imported in R using R tximport package [11] and summarized at the gene level using the "salmon_tx2gene.tsv" file generated by the nf-core pipeline to create a DESeqDataSet object to which a "~group" design combining "Cell.Line" and "Genotype" information was applied.

Comparison of shCTL and shPTBP1 samples was performed in each of the four cell lines (MOLM-14, MV-4, OCI-AML3 and U937) by comparing, for each cell line, the 2 shCTL replicates to the 4 shPTBP1 replicates originating from 2 different shPTBP1 construct (i.e. 2 shPTBP1#1 replicates plus 2 shPTBP1#2 replicates). CombatSeq [12] was used to adjust for batch effects associated with the replicate variable as evidenced by preliminary data analysis using principal component analysis. Log2FoldChange values provided by DESeq::results function were shrunken using DESeq::lfcShrink function with option type = "normal". Changes in gene expression with a p-value adjusted using Benjamini and Hochberg correction (padj )<= 0.05 were considered significant.

**Differential RNA Splicing Analysis**

Differential splicing events between shPTBP1 and shCTL samples were identified using rMATS turbo (v4.1.2) with Maser filtering (F.T. Veiga D (2025). *maser: Mapping Alternative Splicing Events to pRoteins*. R package version 1.29.0, https://bioconductor.org/packages/maser) with the ENSEMBL human genome annotation GRCh38 v109**.** Differential Splicing events were filtered for those with an inclusion level difference (deltaPSI) > 10%, an FDR < 0.05, and average reads >10. Functional consequences of PTBP1 depletion on isoform switching were analyzed using IsoformSwitchAnalyzeR (v1.18.0) and various functional annotation tools (e.g., Pfam, CPAT, SignalP, IUPred2A, Deeploc2, DeepTMHMM).

**Individual Cross-Linking Immunoprecipitation (iCLIP), sequencing and analysis**

PTBP1:RNA interactome in MOLM-14 cells was annotated using individual cross-linking immunoprecipitation (iCLIP). Briefly, cultured MOLM-14 AML cells were washed with ice-cold PBS and UV- irradiated (254 nm, 300 mJ/cm2 using a Stratalinker 2400) prior cell lysing using ice-cold RIPA buffer at 4°C followed by sample sonication (10 seconds cycle, three times). After centrifugation at 15000 rpm, genomic DNA and partial RNA digestion was performed by treating samples with TurboDNase (10 U/ml of lysate, ThermoFisher Scientific) and RNase I (0.167 U/ml, ThermoFisher Scientific) for 3 minutes at 37°C. PTBP1-RNA complexes were immunoprecipitated using 3 μg. of αPTBP1 antibody (mouse IgG1, clone 1, ThermoFisher Scientific) previously coupled to 50 μl of protein G dynabeads (ThermoFisher Scientific). A mouse IgG antibody (clone MOPC-21) was used as negative control. Beads were stringently washed after immunoprecipitation using high-salt washing buffer (50 mM Tris-HCl pH 7.4, 1 M NaCl, 1 mM EDTA, 1% NP-40, 0.1% SDS and 0.5% sodium deoxycholate) and PNK washing buffer (20 mM Tris-HCl pH 7.4, 10 mM MgCl2, 0.2% Tween-20). 3'end RNA dephosphorylation was performed using FastAP alkaline phosphatase (ThermoFisher Scientific) and PNK (New England Biolabs). RNA was linked to a pre-adenylated L3-IR-App adaptor (/5rApp/AG ATC GGA AGA GCG GTT CAG AAA AAA AAA AAA /iAzideN/AA AAA AAA AAA A/3Bio/ coupled with IRdye-800CW-DBCO (LI-COR)) by using T4 RNA ligase I (New England Biolabs) and PNK to visualise protein:RNA complexes. 76. Samples for sequencing were ligated to a non-infrared L3-ATT-App DNA Linker (/5rApp/WN ATT AGA TCG GAA GAG CGG TTC AG/3Bio/) before RNA-protein complexes separation by SDS-PAGE electrophoresis, transfer into a nitrocellulose membrane, and extraction using PK buffer (100 mM This-Cl pH 7.5, 100 mM NaCl, 1 mM EDTA and 0.2 % SDS) with 20 U proteinase K (Roche) at 50°C for 60 minutes. RNA was isolated using phenol/clorophorm extraction and ethanol precipitation, retrotranscribed into cDNA using the SuperScript IV reverse transcriptase (ThermoFisher Scientific) and irCLIP_ddRT_primers sharing a common backbone (/5Phos/WWW-barcode-NNNN GAA TAG GAA GAG CGT CGT GAT/iSp18/GGA TCC/iSp18/TAC TGA ACC GC but with a unique barcode for multiplexing (TCACA, TCCAC or CAGAA). After cDNA purification and circularization with CircLigase II (Epicentre), it was amplified using Solexa P5/P7 primers and sequenced in a DNBseq platform from BGI Genomics (100bp, single-end sequencing).

iCLIP data was mapped to the human genome GRCh38/hg38 and analyzed using the iMaps pipeline (Hallegger M, Cell 2021; Diaz-Munoz M, Nat Commun. 2017) in Flow (https://app.flow.bio/). PTBP1 crosslink sites were filtered for those with an FDR ≤ 0.05 and an extended_score ≥ 3**.** Genomic distribution of PTBP1 crosslink sites was annotated using ChIPSeeker with TxDb.Hsapiens.UCSC.hg38.knownGene annotation.

**Definition of direct PTBP1 targets**

Direct PTBP1 targets were defined as transcripts that were alternatively spliced upon PTBP1 depletion in MOLM-14 and bound by PTBP1 in intronic or exonic regions flanking the alternative splicing site in at least 2 out of 3 iCLIP replicates. More precisely, genes undergoing PTBP1-dependent exon skipping (SE) events were defined as direct PTBP1 targets if any PTBP1 binding event was found to occur either in the intron located upstream the skipped exon, in the intron located downstream the skipped exon or if PTBP1 binding events were both identified in introns upstream and downstream the skipped exon (see Suppl. Figure S5D). Genes undergoing PTBP1-dependent mutual exclusive exons (MXE) inclusion events were defined as direct PTBP1 targets if any PTBP1 binding event was found to occur either in any of the introns located upstream the mutually exclusive exon #1 (*i.e.* MXE intron #1), between mutually exclusive exon #1 and mutually exclusive exon #2 (*i.e.* MXE intron #2), downstream mutually exclusive exon #2 (*i.e.* MXE intron #3) (see Suppl. Figure S5D). Genes undergoing PTBP1-dependent intron retention (RI) events were defined as direct PTBP1 targets if any PTBP1 binding event was found to occur either in the exon located upstream the retained/skipped intron or in the exon located downstream the skipped/retained intron (see Suppl. Figure S5D). Genes undergoing PTBP1-dependent Alternative 3' Splice Site (A3SS) events were defined as direct PTBP1 targets if any PTBP1 binding event was found to occur between A3SS "exon_flanking_end" coordinates and "exon_long_end" coordinates of a A3SS AS event (see Suppl. Figure S5D). Genes undergoing PTBP1-dependent Alternative 5' Splice Site (A5SS) events were defined as direct PTBP1 targets if any PTBP1 binding event was found to occur between "exon_long_start" and "exon_long_end" coordinates of a A5SS AS event (see Suppl. Figure S5D).

**ShRNA sequences**

**Ren.713 (*Renilla Luciferase*, control)**
TGCTGTTGACAGTGAGCGCAGGAATTATAATGCTTATCTATAGTGAAGCCACAGA
TGTA**TAGATAAGCATTATAATTCCTA**TGCCTACTGCCTCGGA

**PTBP1.1206 (#1, human *PTBP1*)**
TGCTGTTGACAGTGAGCGCGCGTGAAGATCCTGTTCAATAGTGAAGCCACAGA
TGTA**TTGAACAGGATCTTCACGCGCT**TGCCTACTGCCTCGGA

**PTBP1.2427 (#2, human *PTBP1*)**
TGCTGTTGACAGTGAGCGCTAGCAAGATGATACAATGGTATAGTGAAGCCACAG
ATGTA**TACCATTGTATCATCTTGCTAT**TGCCTACTGCCTCGGA

#### REAGENT TABLE

| PCR Primers | Manufacturer | Sequence (5'->3') |
|---|---|---|
| CDC42_Fwd | Eurogentec | AGGCTGTCAAGTATGTGGAG |
| CDC42_Rev-v1 | Eurogentec | TCATAGCAGCACACACCTGC |
| CDC42_Rev-v2 | Eurogentec | ACAGAGGTTGCTCTAAGGTG |
| ARHGEF1_Fwd | Eurogentec | CAATGAGCCTGGAGTCCTT |
| ARHGEF1_Rev-v1 | Eurogentec | GAGGCTCTTCTGGTTCAAGC |
| ARHGEF1_Rev-v4 | Eurogentec | AACTGGGGTCCATGTCCA |
| GAPDH_Fwd | Eurogentec | ACCACAGTCCATGCCATCAC |
| GAPDH_Rev | Eurogentec | TCCACCACCCTGTTGCTGTA |
| **WB antibody** | **Manufacturer** | **Reference** |
| ACTIN | Sigma-Aldrich | MAB1501 |
| ACTININ | Cell signaling | 3134 |
| HSP90 | Cell signaling | 4874 |
| PTBP1 | Invitrogen | 32-4800 |
| CDC42 | Cell signaling | 2466 |
| ARHGEF1 | Proteintech | 11363-1-AP |
| PKM1 | Cell signaling | 7067 |
| PKM2 | Cell signaling | 4053 |
| **FACS antibody** | **Manufacturer** | **Reference** |
| mCD45.1-PC5.5 | BD (Becton Dickinson) | 560580 |
| hCD45-APCH7 | BioLegend | 368518 |
| hCD45-APC | BioLegend | 304011 |
| hCD44-FITC | BioLegend | 338804 |
| hCD33-PE | BD (Becton Dickinson) | 555450 |
| AnnexinV-BV421 | BD (Becton Dickinson) | 563973 |
| AnnexinV-APC | BD (Becton Dickinson) | 550474 |
| PI | SIGMA-ALDRICH | P4864 |
| Anti-Puromycin-AF647 | BioLegend | 381508 |
| Precision Count Beads | BioLegend | 424902 |
| **Drugs** | **Manufacturer** | **Reference** |
| Apigenin | MedChemExpress | HY-N1201 |
| CASIN | MedChemExpress | HY-12874 |
| Harringtonine | Combi-Blocks | HD-9179 |
| Venetoclax | MedChemExpress | HY-15531 |
