## Supplementary material for "A PTBP1–CDC42 splicing axis regulates leukemia growth and venetoclax sensitivity in acute myeloid leukemia": Table S1

| Name | FAB | Karyotype | FLT3 | NPM1 | DNMT3A | NRas | KRas | IDH1/2 | CEBPa | p53 | PTEN | Kit | c-Myc |
|---|---|---|---|---|---|---|---|---|---|---|---|---|---|
| MOLM-14 | M5 | complex | ITD | | | | | | | | | | overexpressed |
| MV4-11 | M5 | complex | ITD | | | | | | | | | | overexpressed |
| OCI-AML3 | M4 | complex | | type A | R882C | + | | | | | | | |
| U937 | M5 | t(10;11)(p13;q14) | | | | | | | | deleted | null | | overexpressed |

**Table S1.** FAB classification and genetic features of AML cell lines used for this study.
