## Supplementary material for "A PTBP1–CDC42 splicing axis regulates leukemia growth and venetoclax sensitivity in acute myeloid leukemia": Table S2

| Patient | Karyotype | FLT3 | NPM1 | DNMT3A | NRas | KRas | IDH1/2 | CEBPa | p53 | PTEN | Kit | c-Myc | Others |
|---|---|---|---|---|---|---|---|---|---|---|---|---|---|
| TUH02 | | | mut | | | | | | | | | | |
| TUH88 | | ITD | mut | mut | | | | | | | | | RAD21 |
| TUH90 | | ITD | mut | | | | IDH2m | | | | | | |
| TUH101 | | | | | | | | | mut | | | | |
| TUH104 | | | mut | mut | | mut | IDH1m | | | | | | MGA PTPN11 |
| TUH170 | | | | | mut | | | | | | mut | | CBFB::MYH11 |
| TUH182 | | | | | | | | | | | | | GATA2 RUNX1 |
| TUH211 | | | | | | | | | | | | | PTPN11 |
| TUH248 | | TKD | mut | | | | IDH2m | | | | | | SRSF2 |
| TUH250 | tri7 tri19 tri20 tri4 | | | | | mut | | | | | | | ASXL2 TET2m ZRSR2 |
| TUH254 | | | | | | | | | | | | | |
| TUH278 | in(16) | | | | | | | | | | | | |
| TUH279 | tri8 | | | | | | | | | | | | ALAL BCOR EP300 RIT1 RUNX1 STAG2 TET2 |
| TUH280 | | | mut | | | | | | | | | | ASXL1m TET2m |
| TUH284 | | | mut | mut | | | | | | | | | TET2m |
| TUH287 | | ITD | | | | | | | | | | | ALAL |
| TUH288 | inv(16) | | | | | | | | | | | | |
| TUH291 | | | | mut | | | IDH2m | | | | | | |
| TUH292 | | | mut | mut | | mut | | | | | | | TET2m |
| TUH293 | | ITD | | | | | | | | | | | NUP98::HOXC13 |
| TUH297 | | TKD | mut | mut | | | | | | | | | KAT6B PTPN11 |
| TUH298 | | ITD | | mut | | | | mut | | | | | MLL-PTD |

**Table S2.** Genetic features of patient samples (TUH) used in Figure S1D.
